# *SF3B1* mutation generates novel peptides from long non-coding RNAs

**DOI:** 10.64898/2026.10.15.759235

**Authors:** Supriyo Bhattacharya, Mike Fernandez, Yiming Wu, Meiling Jin, Patrick Pirrotte, Krystine Mansfield, Tae Hyuk Kang, Yuqi Zhao, Nicole Lammana, Xiwei Wu, Lili Wang

## Abstract

While mRNA alternative splicing caused by somatic mutation in *SF3B1* is well-characterized, long non-coding RNAs (lncRNAs) impacted by this mutation are completely unknown. Here, we report long-read sequencing of full-length transcripts from cell lines with and without SF3B1 mutation. We present ALLR (Annotation of human lncRNAs using Long-Read sequencing), a computational workflow to identify and quantify lncRNAs, and to annotate their potential roles in impacting cellular functions. Harnessing matched short-read RNA sequencing and proteomic data, we demonstrate that *SF3B1* mutation generates thousands of novel lncRNAs with diverse levels of alternative splicing. Moreover, *SF3B1* mutation-associated lncRNAs produce translatable peptides with putative functions, immunogenicity, and potential to bind proteins with impact on cellular signalling networks. 47 lncRNAs could also be detected in primary chronic lymphocytic leukemia (CLL) samples and are associated with significant risks towards adverse disease prognosis, in conjunction with other known CLL risk factors. Our work demonstrates the utility of long-read sequencing for lncRNA research and provides a new territory of functional RNA molecules that SF3B1 mutation may use to impact cancer.

**GRAPHICAL ABSTRACT:** 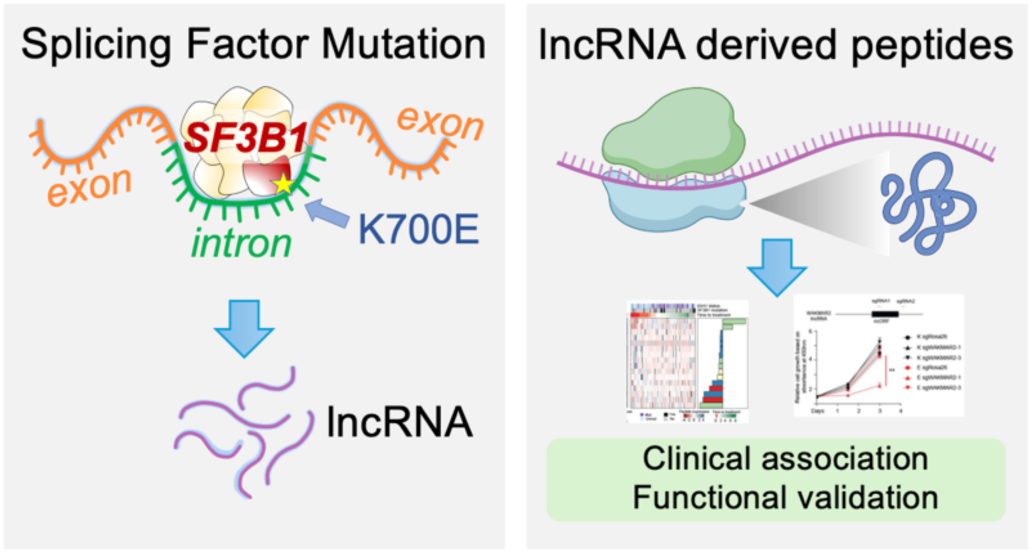

## INTRODUCTION

Long non-coding RNAs (lncRNAs) are a diverse class of RNA molecules species that exceed 200 nucleotides (nt) in length and rarely encod proteins(1,2). These molecules have emerged as an essential class of regulatory molecules for physiological and pathological conditions including cell development and differentiation, and various human diseases such as cancer(1,2). Compared with protein-coding genes, lncRNAs are generally shorter, lower-expressed, less evolutionarily conserved, and exhibit greater tissue-specific expression(1,2). In fact, most of the lncRNAs are still lacking functional annotation, partially due to incomplete and low-quality transcriptome data that limits the accurate identification and structural characterization of these transcripts.

Long-read sequencing opens up a new era of transcriptome annotation(3–5). While previous short-read-based transcriptome reconstruction methods have generated a large number of lncRNA annotations, the quality of software-reconstructed annotations is often doubtful due to the inherent challenges in reconstructing transcript structures from shorter sequence reads. Recently, GENCODE consortium has developed RNA Capture Long Seq (RNA CLS), a method that combines targeted RNA capture with long-read sequencing(6). This approach has significantly increased the annotated complexity of targeted loci, providing a deeper understanding of the genomic features of lncRNAs including their promoters, gene structures, and potential for protein-coding. Thus, long-read sequencing allows the generation of high-quality, full-length transcript models at high-throughput scales.

Alternative RNA splicing gives rise to transcriptomic and proteomic diversity. Human cancers exhibit widespread changes in RNA splicing, which can be caused by somatic mutations or dysregulation of the splicing regulatory machinery(7). *SF3B1,* a splicing factor, is frequently mutated across multiple cancer types including chronic lymphocytic leukemia (CLL), myeloid dysplasia, acute myeloid leukemia, uveal melanoma, breast cancer, and pancreatic cancer(8–11). Previous studies have consistently reported that this gene mutation impacts transcriptome-wide RNA splicing(12–14). However, the impact of *SF3B1* mutations on lncRNA production and their alternative splicing remains poorly understood. Here, we developed a workflow to characterize lncRNAs associated with *SF3B1* mutation by long-read sequencing and to annotate the potential roles of these lncRNAs using *in silico* analysis, providing a new dimension for understanding splicing factor mutations in cancer biology.

## MATERIAL AND METHODS

### Cell lines and culture

HEK293T cells were cultured in DMEM (Gibco) with 10% FBS (Omega Scientific) and 100ug/ml penicillin-streptomycin (Gibco). Nalm-6, HG-3, and MEC-1 cells were cultured in RPMI (Gibco) with 10% FBS (Omega Scientific) and 100ug/ml penicillin-streptomycin (Gibco). Nalm-6 cell lines endogenously expressing *SF3B1* K700K or K700E were purchased from Horizon Discovery. HG-3 and MEC-1 cell lines endogenously expressing *SF3B1* K700K or K700E were engineered as previously described(15). All cell lines were regularly tested and confirmed to be negative for mycoplasma.

### Lentiviral packaging

To package lentivirus, 3μg of the viral plasmid donor, 0.4μg of VSV-G, and 1.5μg psPAX2 were mixed with PEI-MAX 40K (Polyscience) in 1:3 ratio of combined molecular mass in ug to PEI-MAX 40K volume in μl and transfected into seeded HEK293T in 6-well format. Lentiviral supernatant was collected at 72hrs and 96hrs after transfection, pooled, and filtered through a 0.45μm PES syringe filter (Bioland Scientific) before being concentrated with ultracentrifugation at 88,000g for 2hrs at 4°C. To transduce suspension cells, 0.5 million cells were seeded into a 48-well format with 8ug/ml polybrene and concentrated lentivirus and spin-infected at 1000 g for 1.5hrs at 37°C. The media was changed 3hrs after the spin-transduction. Transduced cells were then selected with 2μg/ml puromycin (InvivoGen) for 5 days.

### Flow cytometry and cell proliferation assays

All flow cytometry were performed on a Fortessa X20 (BD Biosciences), and flow data were analyzed using FlowJo version 10.8.2 (BD Biosciences). DAPI was used as a viability dye. For the small-scale essential RNA screening experiment, cells were transduced to ∼40-60% transduction efficiency as measured by RFP expression. Ratios of RFP^+^ to RFP^-^ cells were used to measure cell proliferations. CCK8 assays were also used to measure cell proliferation. Briefly, 20,000 cells were seeded onto 96-well plates in 200ul final media volume and cultured overnight. 100μl of suspended cells were incubated with 10ul of CCK-8 reagent (Dojindo) for 3hrs at 37°C. Absorbance readings were taken using an Infinite M1000 Pro plate reader (Tecan) at 450nm at 24hrs, 48hrs, and 72hrs post-seeding. Cell proliferation results were plotted using Graphpad Prism (Dotmatics).

### Immunoblotting analysis

The antibodies used in this study include: anti-FLAG (#MA1-91878, Invitrogen), GAPDH (#sc-365062, Santa Cruz Biotechnology), goat anti-rabbit IgG secondary antibody (#65-6120, Invitrogen), and goat anti-mouse IgG secondary antibody (#65-6520, Invitrogen). Cells were harvested, pelleted through centrifugation, and washed with PBS buffer. Cell lysates were prepared by lysing cells for 30min on ice in Pierce RIPA buffer (Invitrogen) supplemented with Pierce protease and phosphatase inhibitors (Invitrogen), PMSF (Invitrogen), and sodium orthovanadate (New England Biolabs). Cell lysates were then sonicated and cleared of debris through centrifugation at 12,000 g for 20min at 4°C. Protein quantification was done using a Pierce BCA kit (Invitrogen). Then, 20μg of cell lysate was loaded onto a 4-12% BOLT gel (Invitrogen) and ran at 50V for 30min then 100V for 3hrs in MOPS buffer (Invitrogen). Finally, protein fractions were transferred onto PVDF iBlot 2 membranes (Invitrogen) at 20V for 8min. PVDF membranes were sectioned and then blocked in 5% BSA in 1x TBST for 1hr followed by overnight blotting with primary antibodies. Membranes were washed in 1x TBST the next day, blotted with appropriate secondary antibodies, and then developed in Clarity Max ECL (Bio-Rad). Images were captured on a ChemiDoc (Bio-Rad) and analyzed using Image Lab (Bio-Rad).

### Splice variant qPCR

Total RNA was extracted using Trizol (Ambion) while cDNA was generated using the High-Capacity cDNA Reverse Transcription Kit (Applied Biosystems) per manufacturer’s protocol. 5ng of total cDNA was used as inputs for qPCR on the QuantStudio 12K Flex (Applied Biosystems). OligoDT (Invitrogen) was substituted for random hexamers. To calculate relative ratio of splice variants, The Ct values for the normal and alternatively spliced isoforms were inputted into the following formula: 2^ΔΔCt(Normal)-ΔΔCt(Aberrant)^. The results were graphed using GraphPad Prism (Dotmatics). Primers to detect splice variants are listed in Supplementary Table 7.

### Molecular cloning

All primers used for cloning are listed in Supplementary Table 5. To generate a ZNF609 peptide overexpression plasmid, a DNA fragment (Twist Bioscience) containing a 3xFLAG-ZNF609-ORF sequence was amplified using Q5 polymerase (New England Biolabs) and cloned into pLVX-IRES-ZsGreen (TakaraBio) between SpeI and BamHI (New England Biolabs). For Cas9-related experiments, sgRNAs were designed using CRISPR-Cas9 guide RNA design tool (Integrated DNA Technologies) and cloned into pLKO.5 Puro-2A-RFP using Golden Gate cloning. For CasRx-related experiments, gRNAs were designed using DeepCas13(16) and cloned into pLXR002. All sgRNA sequences are listed in Supplementary Table 6.

### ISO-seq sequencing (ISO-seq)

Total RNA was extracted from Nalm6K and Nalm6E cells using the RNeasy Mini Kit (Qiagen) following the manufacturer’s instructions. Full-length cDNA was synthesized from total RNA using the NEBNext Single Cell/Low Input cDNA Synthesis and Amplification Module (New England Biolabs, Inc.) and Iso-Seq Express Oligo Kit (Pacific Biosciences) according to the PacBio Iso-Seq Express Template Preparation for Sequel and Sequel II Systems protocol. 300 ng of total RNA was primed with NEBNext Single Cell RT Primer Mix and reverse transcribed by NEBNext Single Cell RT Enzyme Mix. After amplifying cDNAs with Iso-Seq Express cDNA PCR Primer, the standard size of the cDNAs (∼2 kb) were chosen by the ProNex Bead (Promega) size-selection. SMRTbell libraries were prepared by ligating hairpin adapters to double-stranded cDNAs using the SMRTbell Express Template Prep Kit 2.0 (Pacific Biosciences). Prepared SMRTbell libraries were sequenced on the PacBio Sequel platform. Libraries were loaded onto a SMRT Cell 1M and sequenced in movie mode with a 20-hour collection time to maximize read length. The primary analyses, including real-time signal processing and base calling were processed by a built-in PacBio Blade Center through Sequel ICS, and result stream directly to SMRT Link (v8.0.0.80529). Sequencing was performed at the integrated genomic core at City of Hope Medical Center.

### Ribosomal profiling (Ribo-seq)

Nalm6 K and E cells were harvested at ∼80% confluency and treated with cycloheximide (100 µg/mL, 5 min at 37 °C) to arrest translating ribosomes. Cells were then washed with ice-cold PBS containing cycloheximide and lysed in lysis buffer (20 mM Tris-HCl pH 7.4, 150 mM NaCl, 5 mM MgCl₂, 1% Triton X-100, 100 µg/mL cycloheximide, and RNase inhibitor). Lysates were clarified by centrifugation at 20,000 × g for 10 min at 4 °C and then treated with RNase I (Ambion) at 25 °C for 45 min to digest unprotected RNA. Digested RNA was mixed with 2× RNA loading dye and resolved on a 15% TBE-urea polyacrylamide gel (one well reserved for a small-RNA ladder). Following SYBR Gold staining, the 26–34 nt region was excised with minimal UV exposure to obtain ribosome-protected fragments (RPFs). Gel slices were crushed and eluted overnight at 4 °C in 300 mM sodium acetate (pH 5.5), 1 mM EDTA, 0.1% SDS. The eluate was passed through a 0.22 µm filter, ethanol-precipitated and pelleted, washed with 75% ethanol, air-dried, and resuspended in nuclease-free water to obtain RPFs. Libraries were prepared using the TruSeq Small RNA Sample Preparation Kit (Illumina) according to the manufacturer’s protocol, with modifications for RPFs. Libraries were quantified with a Bioanalyzer (Agilent Technologies) and sequenced on an Illumina NovaSeq platform to generate single-end 50 nt reads.

### Tandem mass tag Mass spectrometry (TMT MS)

Nalm6K and E cells were lysed with TEAB buffer supplemented with protease inhibitor and PMSF. Lysates (300 μg) were precipitated and digested to obtain peptides. TMT 10-plex labeling was performed according to the manual from Thermo Fisher Scientific, and peptides were fractionated via BPRP HPLC. An 1100 pump (Agilent) equipped with a degasser, and a photodiode array (PDA) detector (Thermo Fisher Scientific) was used. Peptides were subjected to a linear gradient from 3% to 25% acetonitrile in 0.125% formic acid using an Agilent 300 Extend-C18 column (Agilent) and were fractionated into a total of 96 fractions. Mass spectrometry was performed using an Orbitrap Fusion mass spectrometer (Thermo Fisher Scientific) coupled to a Proxeon EASY-nLC 1000 liquid chromatography (LC) pump (Thermo Fisher Scientific). Peptides were detected (MS1) and quantified (MS3) in the Orbitrap. Data analysis was performed using the MAXQUANT software suite. MS2 spectra were searched using the Andromeda search engine against a Uniprot composite database derived from the human proteome containing its reversed complement and known contaminants, along with predicted amino acid sequences of lncRNA derived peptides. Peptide spectral matches were filtered to a 1% false discovery rate (FDR) using the target–decoy strategy.

### Processing of ISO-seq reads

Starting from the raw fastq reads, the FLAIR pipeline (version 1.4)(17) was used to align the reads to the human genome (Flair align, GRCh38 without ALT contigs, as explained in the FLAIR documentation), followed by correction of splice junctions using short reads (Flair correct, please see section: Short reads-based quantification of isoform expression) and reconstruction of full-length RNA isoforms (Flair collapse).

### Processing of Ribo-seq reads

Ribosome profiling (Ribo-seq) reads were first aligned to ribosomal RNA (rRNA) reference sequences using Bowtie to remove rRNA-derived reads (18). rRNA reference sequences were obtained from Ensembl version 95(19). Reads that did not map to rRNA were subsequently aligned to the human reference genome (GRCh38/hg38) using HISAT2 with default parameters (20). Reads with multiple genomic alignments were excluded from downstream analyses. Quality control analyses, including read length distribution, reading frame distribution, three-nucleotide periodicity, and P-site (peptidyl-site) position estimation for ribosome-protected fragments (RPFs), were performed using the R package riboWaltz (21). Only RPFs with lengths between 28 and 34 nucleotides were retained for further analysis. Ribosome occupancy across candidate open reading frames (ORFs) was quantified using Ribo-seq data by calculating the fraction of each ORF covered by at least 10 Ribo-seq reads.

### Short read-based quantification of isoform expression

The ISO-Seq derived transcripts were quantified among the short-read data from Nalm6 and CLL cell lines using SALMON(22). A custom transcriptome fasta was prepared by concatenating the human genome with the transcriptome fasta from Iso-Seq (as described in the SALMON documentation). The short reads from the Nalm6 (K700K, K700E)(23) and CLL (HG3 and MEC1)(15) samples were then aligned to the custom transcriptome to quantify the transcript expressions. Differential expression analysis was performed using DESeq2(23,24). Additionally, differential transcript usage was analyzed using DRIMseq(25).

### Splicing event analysis

Alternative splicing events were detected using Miso(26) employing a splice events database from ISO-Seq transcripts and aligned short reads for quantification. In brief, a custom splice events database was generated through rnaseqlib(27), using the Iso-Seq derived transcripts as input and the flanking-rule parameter set to ‘commonshortest’. Using this database, alternative splicing events in five separate categories (A3’SS, A5’SS, SE, MXE and RI) were quantified in Miso, followed by comparison between K700K and E samples. Events with a bayes factor > 5 were defined as significantly different between K700K and E. Splice junctions and the corresponding read counts were obtained from aligned short reads using STAR v 2.6 (28). Differential junction usage was analyzed using DEXseq (29). For comparison with SJs from RJunBase, the annotation data downloaded from http://www.rjunbase.org/rJunBase/download/toIndex was imported into R as a GRanges object using the GenomicRanges package (30). SJs from ISO-Seq were matched with those from RJunBase using the findOverlaps() function from GenomicRanges with the criteria that both ends of the junction should match with a maximum gap of 3 nt. Cryptic splice site DNA motifs were plotted using the ggseqlogo package(31).

### DIA-MS data from CLL patients

The raw DIA-MS data from 117 CLL patients were downloaded from the PRIDE databank (PRIDE ID: PXD022198)(32). Spectra were searched against a protein FASTA of 806 lncRNA derived peptides using Spectronaut v18.0 with default settings(33). In summary, trypsin theoretical digestion was applied, allowing for up to 2 missed cleavages. Peptides were filtered for those with length 4-52 amino acids. Both proteins and peptides were filtered for FDR < 1%. Methionine oxidation and N-terminal acetylation were set as dynamic modifications with cysteine carbamidomethylation as a static modification. Peptide abundances were normalized using variance stabilizing normalization (vsn package(34)) in R v3.6(35).

To analyze the potential contribution of the ncORFs towards disease complication (progression free survival), we first performed a univariate regression analysis of log-normalized ncORF expressions along with patient clinical and genomic attributes using progression free survival as the dependent variable (metadata entry:‘Time to treatment’). Multivariate regression was performed on each ncORF using the log-normalized expression as independent variable and the time to treatment as dependent variable, along with key clinical and genomic attributes (i.e., IGHV status, *del*(11q), *del*(17p), trisomy 12, mutations in *SF3B1*, *TP53*, *ATM*) as covariates. Univariate and multivariate analyses were conducted using the R package ‘survivalAnalysis’.

## RESULTS

### ALLR (Annotation of human lncRNAs using long-reads sequencing), functional annotates novel lncRNAs from long-read sequencing

We have developed a parallelizable computational pipeline to identify and functionally Annotate human LncRNAs using Long-Read sequencing approaches (ALLR) from PacBio Iso-Seq and Nanopore sequencing (**Fig. 1A**). Although targeted primarily towards long-read sequencing, ALLR can also analyze lncRNAs assembled from short-read RNA sequencing (RNA-seq) data. ALLR addresses the following aspects of the detected lncRNA transcriptome: (1) expression quantification and optional comparison between biological conditions; (2) discovery of novel lncRNAs by comparing with existing lncRNA databases (e.g. NONCODE); (3) identification of disease-associated lncRNAs; (4) quantification of lncRNA-associated splicing events; (5) detection of translatable lncRNAs using mass spectrometry/ribosome profiling (Ribo-seq) and functional characterization of resulting protein products; (6) functional analysis of lncRNA-protein interaction.

**Fig. 1.**
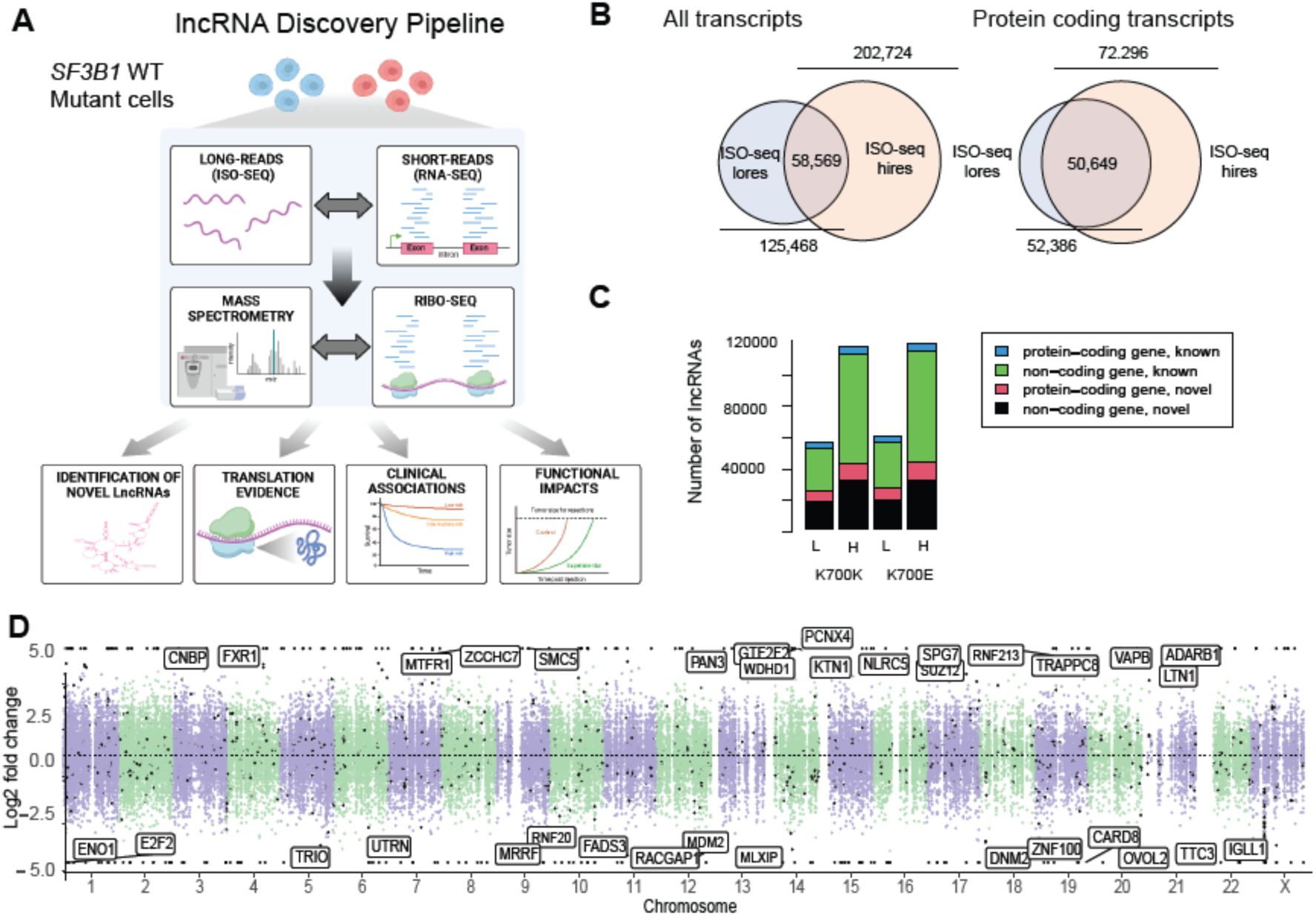
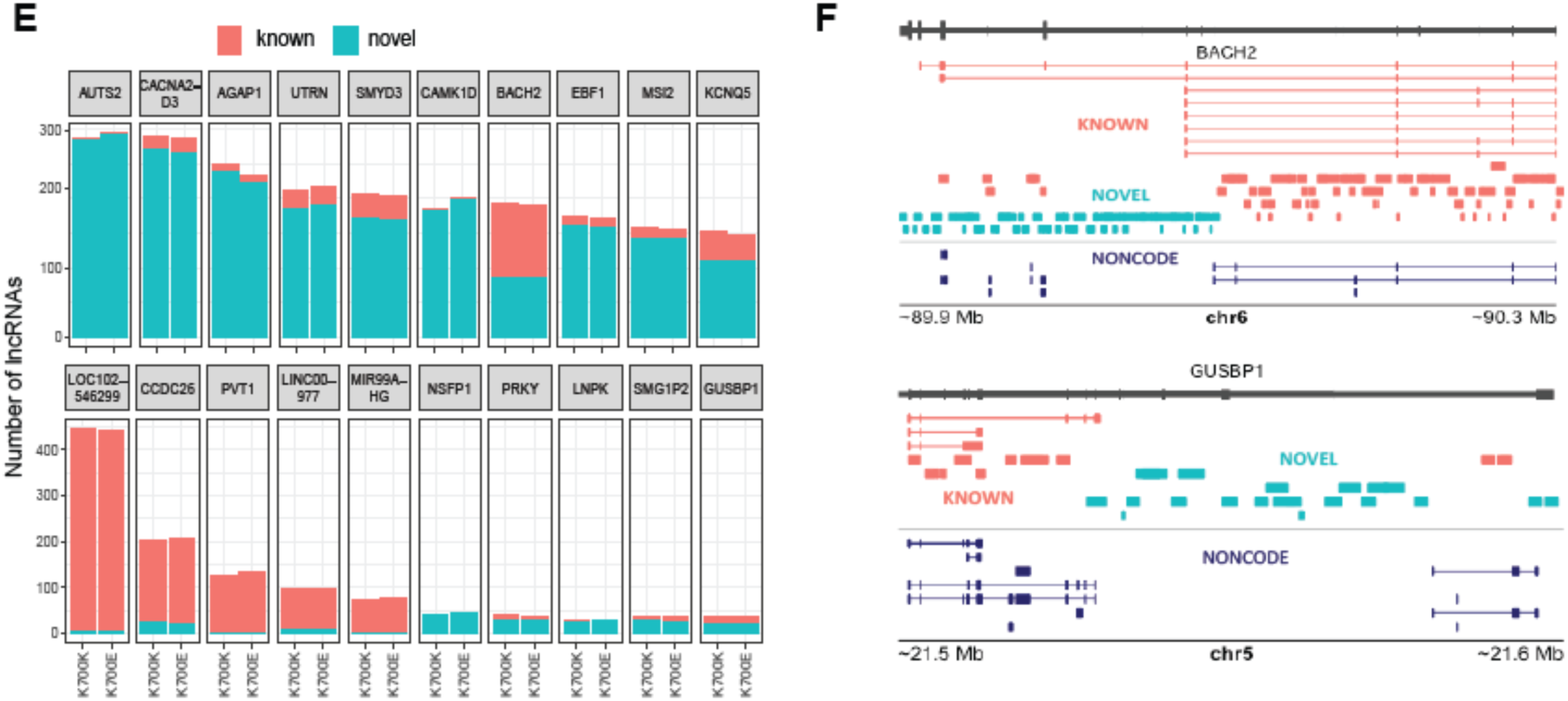
*SF3B1* mutation impacts lncRNA landscape. **(A)** Schematic of the lncRNA discovery pipeline, **(B)** Overlap of identified transcripts between ISO-Seq^lores^ and ISO-Seq^hires^. **(C)** Number of known and novel lncRNAs identified from protein-coding and non-coding genes in the K700K and E samples from ISO-Seq^lores^ (L) and ISO-Seq^hires^ (H). **(D)** Manhattan plot showing genome-wide distribution of lncRNAs identified using ISO-Seq^hires^. X-axis represents the genomic coordinates for each lncRNA and the Y-axis the corresponding log fold change (K700E vs. K700k) as estimated using short-reads RNA-seq. LncRNAs showing statistically significant expression differences between K700K and E (FDR < 0.1) are highlighted as black dots. The top differentially expressed lncRNAs in each chromosome are labeled. **(E)** Top 10 protein-coding genes with the highest number of known and novel lncRNAs showing the dedicated lncRNA genes. **(F)** Examples of lncRNAs from one protein-coding and one non-coding gene. The known and novel transcripts are colored orange and green respectively. The lncRNAs annotated in the NONCODE database are shown in dark blue.

Starting with raw fastq reads from long-read sequencing data, ALLR uses FLAIR(17) to align the reads to the genome and constructs full length isoforms with high accuracy. Coupled with short-read RNA seq data, FLAIR also quantifies the isoform expression (SALMON)(22) and splice variants (MISO)(26) with increased quantification accuracy due to improved sequencing depth. Differentially expressed lncRNAs between biological conditions are then detected using DESeq2(25,36). Moreover, ALLR can distinguish between differential expression of lncRNAs caused by altered splicing, as opposed to expression changes of the parental gene, using DRIMseq and StageR(25).

ALLR analyzes genome-wide lncRNA splicing events with splicing event categories (e.g. intron retention) annotated. It also categorizes the identified lncRNAs as known and novel by comparing against the NONCODE lncRNA database (downloaded in June, 2020). When a lncRNA shows less than 60% genomic overlap with any known lncRNA in the database, it is classified as a novel lncRNA. To identify translatable lncRNAs, the open reading frame (ORFs) originated from the transcript sequences are searched for ribosomal engagement and matched against tandem mass tag mass spectrometric (TMT-MS) spectra for evidence of translated peptide hits (more details in the methods section), that do not resemble any known protein sequences. The detected lncRNA derived peptide sequences are then analyzed for statistical characteristics such as amino acid and predicted secondary structure distributions as well as presence of putative functional motifs(37). Further, we have used the deep learning driven algorithm AlphaFold V2 to predict the tertiary structures of these peptides(38). A significant fraction of these peptides folded into protein-like structures, suggesting that lncRNA-derived peptides could have functional consequences. Lastly, the lncRNA-protein interactions were predicted using the software AnnoLnc 2.0(39), followed by functional characterization of the lncRNAs based on the proteins they interact with. We also assessed the immunogenicity of the lncRNA-derived peptides using the deep learning-based method DeepImmuno(40) and examined their expression and clinical associations in an independent human CLL proteomic dataset. Furthermore, we validated that translatable lncRNAs contribute to *SF3B1* mutation-dependent cell growth.

### *SF3B1* mutation generates novel lncRNAs

With ALLR, we characterized lncRNAs associated with *SF3B1* K700E mutation, one of the most common hotspot mutations on this gene found in cancer. We used isogenic Nalm6 cell lines: Nalm6K700K (Nalm6K) and Nalm6K700E (Nalm6E), for which the cells carry a silent mutation or heterozygous *SF3B1-*K700E mutation at the endogenous locus(23). Two versions of long-read sequencing platform from PacBio were used, SEQUEL II and IIe, referred to as ISO-Seq^lowres^ and ISO-Seq^hires^ respectively. We took this opportunity to assess the efficiency from generational improvements to long-reads technology. While ISO-Seq^lores^ identified about 125,000 isoforms, ISO-seq^hires^ identified about 203,000 isoforms, confirming higher detection power of the newer technology (**Fig. 1B**). About 29% of the ISO-Seq^hires^ isoforms overlapped with those from ISO-Seq^lores^. Among the ISO-Seq^lowres^ isoforms, 55% were protein-coding, while the rest (45%) were noncoding (Supplemental Table S1). In contrast, 64% of the ISO-Seq^hires^ isoforms were noncoding (**Fig 1B**). Among the protein-coding isoforms, 70% were common between ISO-Seq^hires^ and ISO-Seq^lores^. This shows that the increased sensitivity of the newer PacBio technology is advantageous towards the detection of lncRNAs, which are typically of lower expression than protein-coding isoforms(41).

Among the 130,428 non-coding RNAs found in Nalm6K and Nalm6E cells, 130,391 (99.9%) are longer than 200nt, suggesting they are lncRNAs(1,2). By comparing to NONCODE containing 173,112 annotated human lncRNAs(42–46), ISO-Seq^hires^ retrieved 82,992 novel lncRNAs, that have less than 60% overlap with any known lncRNA in the database (**Fig. 1C**). When comparing the expression of differentially expressed lncRNAs between Nalm6K and Nalm6E identified from short-read RNA-seq, 440 and 518 were found to be up- and down-regulated in Nalm6E cells, respectively (FDR < 0.1, **Fig 1D**). While many of these lncRNAs originate from protein coding genes, some are also from known lncRNA genes such as LINC01138, LINC00630, LINC00674, LINC00847, LOC646588 (up in K700E) and LOC100129697 (down in K700E). Notably, several of these lncRNAs were reported to have major roles in several cancers such as LINC01138 (hepatocellular carcinoma(47)), LINC00630 (colorectal(48) and non-small cell lung cancer(49)), LINC00674 (hepatocellular carcinoma(50)) and LINC00847 (non-small cell lung cancer(51), squamous cell carcinoma(52) and hepatocellular carcinoma(53)). We found certain protein-coding and dedicated lncRNA genes to be the sources of multiple known and novel lncRNA isoforms (**Fig. 1E**), with *BACH2* and *GUSBP1* shown as examples (**Fig. 1F**). Taken together, long-read technology in conjunction with efficient in-silico pipelines enables the discovery of multiple novel lncRNAs with potential biological significance.

### Novel splicing events originating from *SF3B1* mutation discovered through ISO-Seq

One of the advantages of long-read sequencing is the identification of alternative spliced isoforms and novel splicing junctions. Using ISO-Seq^hires^ data coupled with the rnaseqlib pipeline(27) (https://rnaseqlib.readthedocs.io), we detected the usage of *de novo* splice junctions (9545) and known junctions from the UCSC Genome Browser repository (130,207) (**Fig. 2A**). Apart from GENCODE, we compared the ISO-Seq-derived splice junctions with the RJunBase database, which catalogs RNA splice junctions identified in both healthy and cancerous human tissues(54). We found 8075 splice junction matches from RJunBase originating from both protein-coding and noncoding isoforms (**Fig. 2A**). We then quantified alternative splicing events at these junctions from the short-read RNA-seq data using MISO(26). We classified five categories of alternative RNA splicing events (alternative 3’ splicing (A3’SS), alternative 5’ splicing (A5’SS), skipped exon (SE), mutually exclusive exons (MXE), and intron retention (IR)). In each category, long-read based splicing junctions showed higher number of events compared to the known junctions from UCSC with the largest difference in the IR events, where long-read discovered 33,048 events compared to known junctions (4217 events) (Supplementary tables 2-3). Also, we detected significantly lower number of splicing events using the *de novo* splicing junctions detected from ISO-Seq^lowres^ compared to ISO-Seq^hires^ (Supplementary table 4). Overall, combining isoform information from long-read sequencing with junction reads from short-read sequencing significantly increases the possibility of discovering novel splicing events due to splicing factor mutation.

**Fig. 2.**
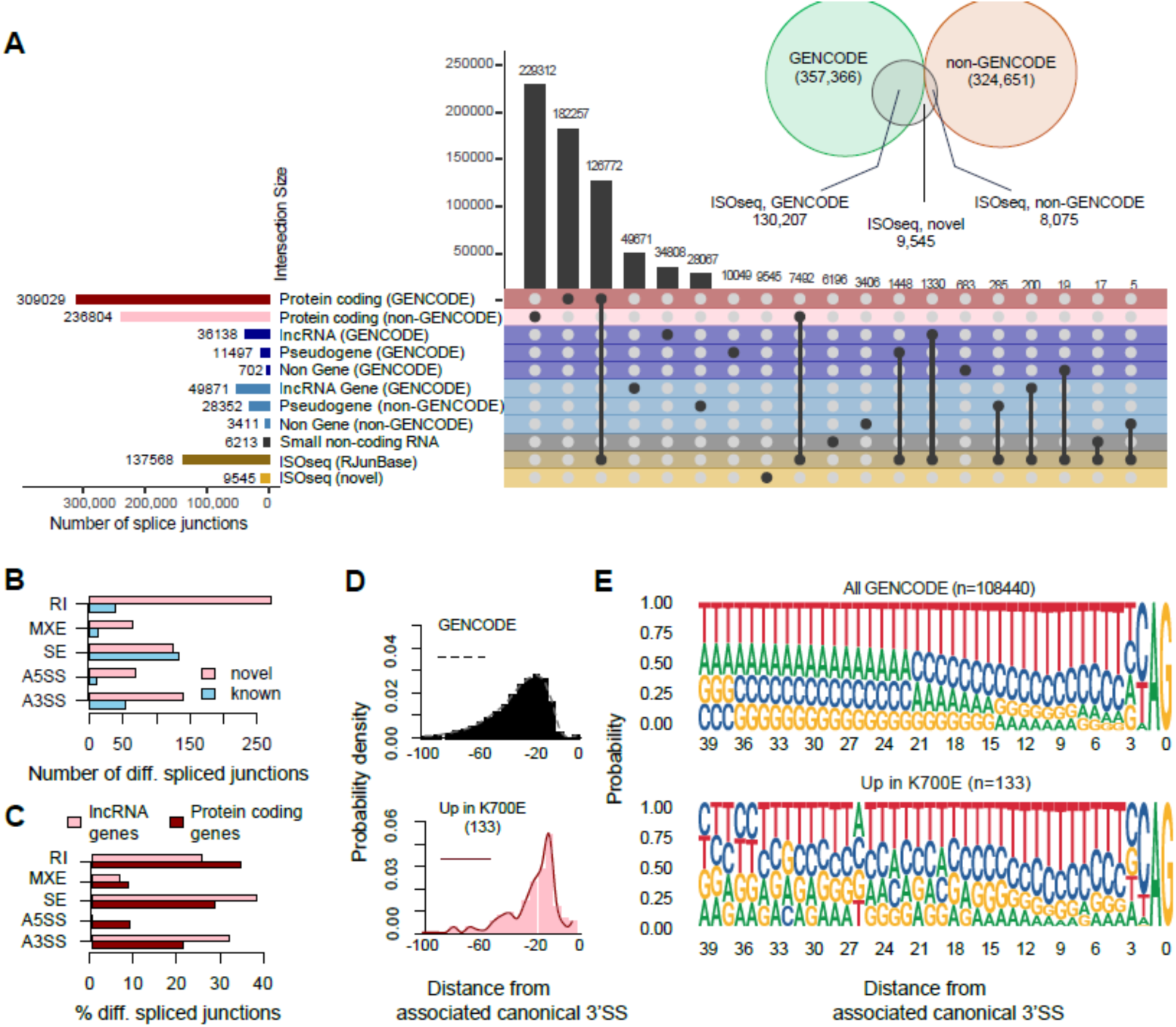
*SF3B1* mutation associated lncRNAs have a splicing pattern similar to mRNA. **(A)** Overview of the SJs identified by ISO-Seq^hires^ in Nalm6 K700K and E samples. Horizontal bars in the Upset plot give the number of SJs belonging to different categories annotated in RJunBase, as well as the ones identified by ISO-Seq^hires^. The row “ISO-Seq (RJunBase)” indicates SJs common between ISO-Seq and RJunBase, while “ISO-Seq (novel)” indicates novel SJs not included in RJunBase. Overlaps between ISO-Seq and each category of SJs in RJunBase are quantified by the vertical black bars. Venndiagram depicts the overlap among ISO-Seq, GENCODE and non-GENCODE SJs from RJunBase. **(B-C)** Number of differentially expressed AS events in K700E vs K in different categories separated into (B) novel and GENCODE annotated, and (C) associated with lncRNA and protein coding genes. **(D)** Distance of the nearest AG (not part of a GAG primer) from the associated canonical 3’SS among all GENCODE annotated SJs (top), distance of cryptic 3’SS from the associated canonical SS, that are exclusively upregulated in K700E (down). **(E)** Enriched nucleotide motifs within 40 nt from the canonical 3’SS among (top) all GENCODE SJs and (down) cryptic 3’SS that are upregulated in K700E.

We further identified both novel and known splicing events associated with *SF3B1* K700E and found a higher number of novel spliced events in all categories except for SE (**Fig. 2B**). Additionally, we found that protein-coding and lncRNA-associated events were similarly distributed among the five splicing event categories except RI and A5’SS events (**Fig. 2C**). Based on the summary of genome-wise distribution of differential splicing events using circular plot (**Fig. S1A**), we discovered that chromosomes 1, 3 and 17 showed higher number of splicing events compared to the rest. Also, there were differences among the proportions of splicing types for each chromosome. For example, chromosomes 11 and 16 showed high proportion of retained introns only, whereas chromosome 18 showed both high A’SS and retained intron splicing events, while chromosome 19 showed higher number of skipped exons. Some chromosomes such as 1 and 17 showed similar number of A3’SS, SE and IR events. The comparison between upregulated and downregulated splicing events indicates no chromosome specific bias as both types of events appear to be randomly distributed throughout the genome.

One of the hallmarks of *SF3B1* mutation is the preferential selection of cryptic 3’ splice site over canonical one(13,23,55,56). To verify whether the ISO-Seq-derived splice junctions reflect this trend, we quantified the splice junction expressions using short-read data (details in methods) and measured the distances of cryptic splice sites that were upregulated in *SF3B1* K700E cells (133 sites, log fold change > 0.1) from their corresponding canonical splice sites. The upregulated splice sites in *SF3B1*K700E showed a narrow distance distribution peaking around 12-24 nt from the canonical splice site (**Fig. 2D**). In contrast, the distance of the nearest AG (non-GAG trimer) upstream of canonical splice sites from GENCODE showed a broad distribution indicating lack of specificity (**Fig. 2D**). Further, motif analysis showed predominance of thymines and cytosines immediately upstream of the canonical AG (indicative of polypyrimidine tract) and an enrichment of adenine about 26 nt from the canonical AG (indicative of branch point selection) (**Fig. 2E**). These are known hallmarks of *SF3B1* mutation reported by multiple groups in the past including our own(13,23,55,56), and validates the *SF3B1* K700E transcriptome revealed by ISO-Seq^hires^.

### ISO-seq identifies multiple disease-associated splice junctions and lncRNAs upregulated in *SF3B1* mutant cells

Using ISO-seq, we identified thousands of novel splice junctions (SJs) that are not annotated in databases such as GENCODE (**Fig. 2A**). To find out whether any of these SJs are relevant to diseases, we queried RJunBase(54) and found 272 SJs that are reported to be exclusively expressed in tumor tissues and not normal healthy tissues (**Fig. 3A**). Of note, the highest number of expressed tumor-specific SJs were common with leukemic tissue, indicating that ISO-seq derived SJs carry a leukemogenic signature reflecting the source of the Nalm6 cell-line. Besides this, we also found SJs expressed in several solid tumor types (**Fig. 3B**). Further, 17 tumor-specific SJs were found to be differentially expressed between *SF3B1* K700E and K700K cells based on short-read sequencing (**Fig. 3C**, p < 0.05). Each of these differentially expressed SJs were shared by multiple isoforms (**Fig. 3D**). ISO-seq enables in-depth analysis of these tumor-specific SJs by examining their positions within the context of full-length isoforms. As an example, one of the tumor-specific SJs (located along the negative strand on chromosome 2) occurs on gene IMMT (**Fig. 3E-F**) and is an alternative 3’ splicing event upregulated in *SF3B1*K700E cells (**Fig. 3F**). Taken together, long-read sequencing, in conjunction with short-read based quantification, enables the discovery of novel SJs that are significantly modulated by splicing defects, potentially leading to aberrant lncRNA expressions.

**Fig. 3.**
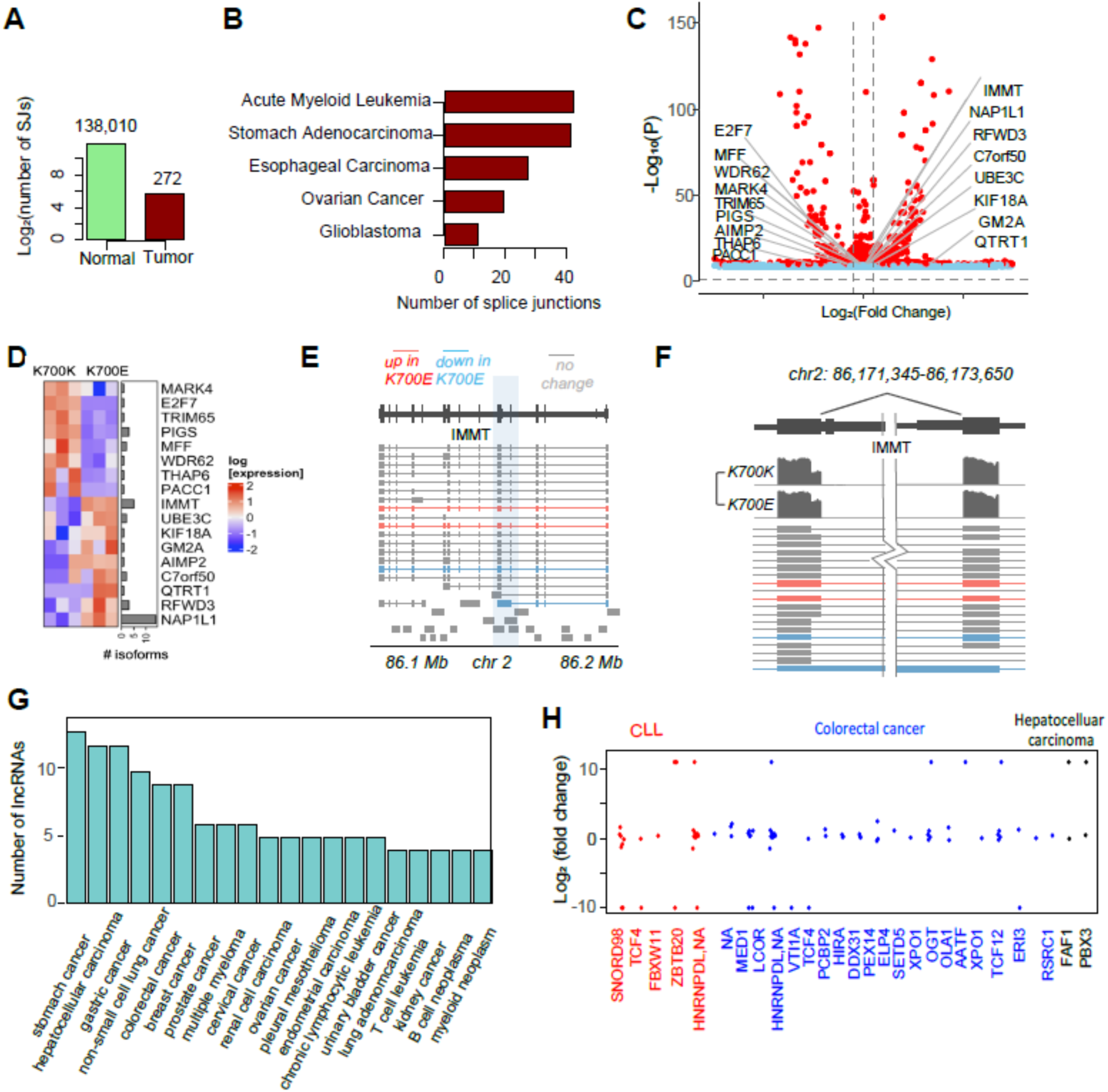
*SF3B1* mutation associated lncRNAs can be found in various disease. **(A)** Number of SJs reported by ISO-Seq^hires^, categorized into normal and tumor specific, based on annotation in RJunBase. Normal refers to SJs that were detected in normal tissue and tumor specific are the ones, exclusively found in tumor samples. **(B)** Top five cancer types among the tumor specific SJs from ISO-Seq^hires^. **(C)** Volcano-plot of SJs found in ISO-Seq^hires^, showing the log fold change in usage between K700E and K, obtained from short reads RNA sequencing. Differentially expressed tumor specific SJs are labeled (*p < 0.05*). **(D)** Expression of the tumor specific SJs labeled in (C) among the replicates of K700K and E obtained from RNA-seq. Adjoining barplot gives the number of isoforms for each SJ from ISO-Seq^hires^. **(E)** Example of a differentially expressed tumor specific alternative 3’ spliced SJ (highlighted by blue shaded rectangle) identified by ISO-Seq^hires^, shown in a genomic diagram. Shown are all isoforms from the parent gene given by long reads sequencing, color coded by their differential expression status. **(F)** expanded view of the tumor specific SJ from (E), along with normalized read densities in K700K and E from short reads sequencing. **(G)** Barplot representing the number of disease associated lncRNAs identified by ISO-Seq^hires^, based on annotation in NONCODE. **(H)** lncRNAs from ISO-Seq^hires^ that overlap with the UCR derived lncRNAs reported by Calin et al., categorized by disease type. lncRNAs uniquely detected in either K700K or K700E (LFC = ±∝) were assigned a LFC of ±10 for displaying in the graph.

To assess the disease relevance of the identified lncRNAs from ISO-seq, we searched the NONCODE database for lncRNAs associated with human cancers. We found 1531 lncRNAs belonging to 166 unique genes from ISO-seq^hires^ with disease association. Next, we specifically examined the lncRNAs that were upregulated in K700E cells (FDR < 0.1). Out of 576 upregulated lncRNAs, 39 were found to be implicated in various diseases (**Fig. 3G**). The diseases representing the highest number of lncRNAs that are upregulated in K700E include stomach cancer, hepatocellular carcinoma, gastric cancer, colorectal cancer, non-small cell lung cancer and breast cancer. The source genes for these lncRNAs include *FTX, NBR2, TP53, ZFAS1, SNHG5, HEIH, INTS7* and *GAS5*.

We also investigated lncRNAs detected in ISO-seq that originate from ultra-conserved regions (stretches of DNA—typically 200 base pairs or longer—that are 100% identical across distantly related species, UCR) in the genome. 88 UCRs associated lncRNAs have been reported to be associated with cancer progression with inhibition of apoptosis(57). When overlapping our lncRNAs from ISO-seq^hires^ transcriptome, we found 27 hits (**Fig. 3H**). The UCR derived lncRNAs showed both up and downregulation in K700E cells. Moreover, they were originally reported in tumor tissue from three human cancers: CLL, colorectal cancer and hepatocellular carcinoma, highlighting that long-read sequencing can be valuable in discovering disease relevant lncRNAs. As an example of a disease relevant lncRNA gene that is implicated in leukemia, we examined *DLEU2* (deleted in lymphocytic leukemia 2), a non-coding gene that produces several microRNAs (Mir-15a, miR-16-1, Mir-455) with tumor suppressor effect(58). The chromosomal region corresponding to the *DLEU2* gene (13q14.3) is deleted in nearly 55% of CLL patients(59). However, this gene is also known to produce several alternatively spliced lncRNAs that are positively correlated with certain cancers(60,61). Of note, a *DLEU2* generated lncRNA ALT1 was shown to promote cell proliferation by acting through the ACE2/Cyclin D1 pathway(62). In total, we found 57 different lncRNAs detected in ISO-seq from the gene *DLEU2* (**Fig. S1B**). These isoforms include both multi-exon transcripts, as well as intronic fragments. Three of these isoforms were significantly upregulated in K700E, according to the short-read RNA-seq data (highlighted in green in **Fig. S1B**). Interestingly, we found a lncRNA isoform unique to K700E that is closely related to ALT1 (highlighted by red box in **Fig. S1B**). The genomic region encoding the tumor suppressor microRNAs Mir-15a/Mir-16-1 is notably transcriptionally inactive, as evidenced by the lack of RNA-seq read density. Taken together, this indicates that that *SF3B1* mutations may modulate lncRNA output from a CLL disease-associated locus.

### Integration of mass spectrometry with ISO-Seq leads to large-scale discovery of translatable lncRNAs with potential mini protein-like folds

Previous works have shown that many lncRNAs are translatable. However, previous evidence primarily relied on a combination of short-read sequencing, ribosomal profiling (ribo-seq), and mass spectrometric (MS) analysis. Due to the absence of reliable technology for sequencing full-length transcripts, the number of detected protein products derived from lncRNAs has remained limited (< 100)(63),(64). By leveraging our long-read data with full-length isoforms annotation combined with MS spectra from the same cells, we identified hundreds of translatable lncRNAs, significantly expanding our knowledge of translatable lncRNA (**Fig. 4A**). We first probed our ISO-Seq derived 130,428 lncRNA sequences for the existence of ORFs encoding for potential peptides. To our excitement, we discovered nearly 2.4 million ORFs greater than 20 amino acids in length. Next, we utilized ribo-seq to assess ribosomal read coverage across these ORFs, measuring the fraction of each ORF covered by at least 10 ribo-seq reads. Strikingly, we identified approximately 54,000 ORFs with more than 50% ribosomal coverage, strongly suggesting potential translation (**Fig. S2A**). We also calculated the conservation of these ORF sequences within the mammalian family using the Phylogenetic Analysis with Space/Time models conservation (PhastCons) method(65). When plotting the Ribo-seq read density against the PhastCons score, we observed that highly conserved ORFs (PhastCons score > 0.8) showed higher ribosomal engagement compared to weakly conserved ORFs (PhastCons score < 0.2) indicating possible evolutionary selection for translation of lncRNA-derived ORFs (p < 2.2×10^-16^) (**Fig. S2B-C**).

**Fig. 4.**
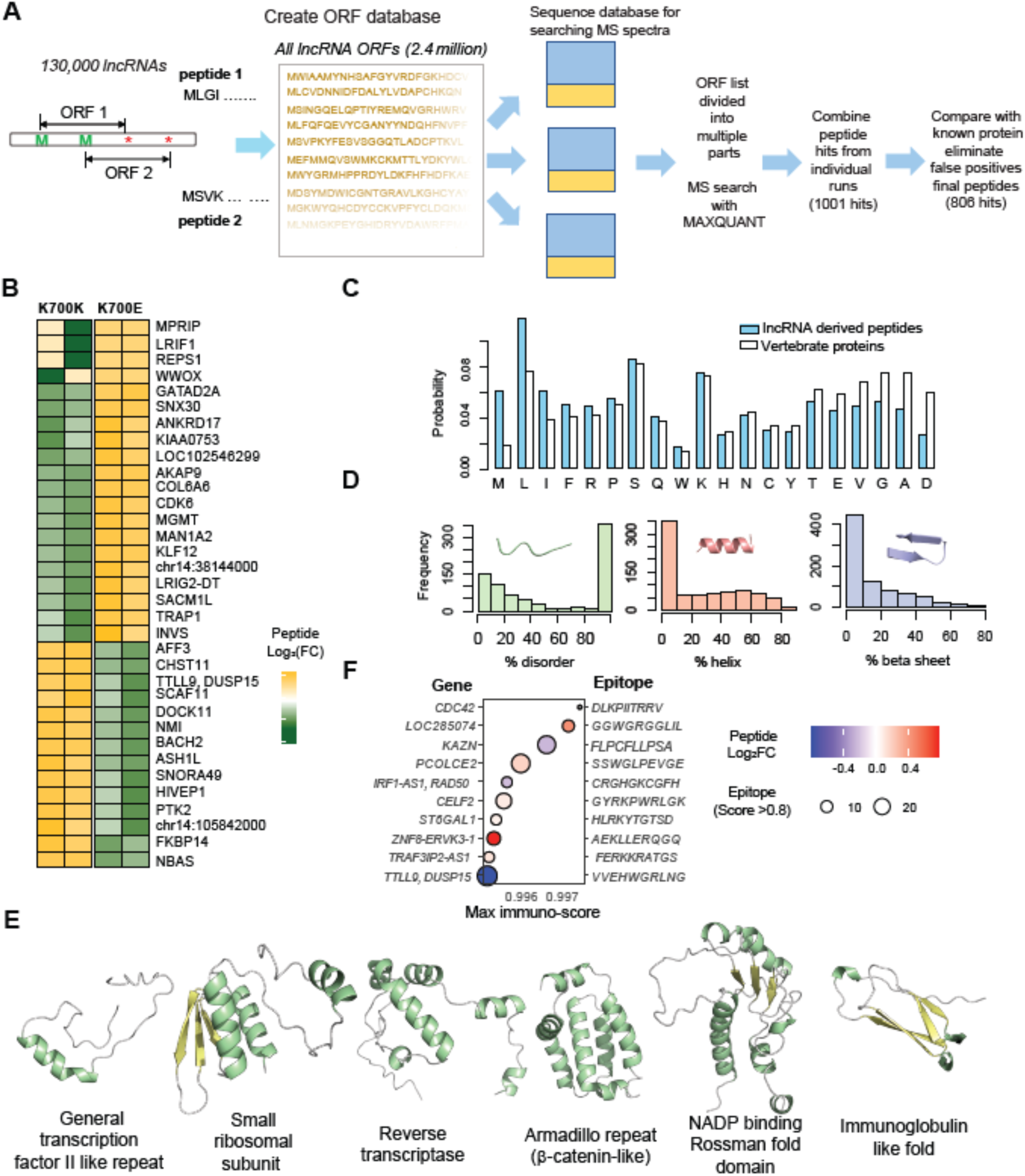
*SF3B1* mutation associated lncRNAs can be translated. **(A)** Pipeline for discovering translated peptides from lncRNA ORFs by integrating MS spectra with long reads isoform information. **(B)** Normalized expressions of differentially expressed lncRNA derived peptides (E vs K, p value < 0.05) in K700K and E (two replicates each) from mass spectrometry. **(C)** Propensities of all 20 amino acids among detected lncRNA peptides and comparison with vertebrate proteins. **(D)** Distribution of secondary structures content among translatable lncRNA peptides, (prediction: RaptorX). Histograms represent the amounts of disordered, helical and beta sheet structures in each peptide sequence as percentage of amino acid length. **(E)** Predicted immunogenicity of the translatable lncRNA peptides using DeepImmuno; Highlighted are peptides with highest immunogenic epitopes (epitope sequence shown on the right) according to the predicted immunogenicity score; dots are colored based on the log fold change (E vs K) from mass spectrometry and the dot sizes are according to the number of immunogenic epitopes within the ORF (score > 0.8). **(F)** Examples of folded structures of lncRNA derived peptides, as predicted by AlphaFold 2.0 and the associated functional domains identified by InterPro.

Building on these findings, we conducted a systematic search of the 2.4 million lncRNA ORFs for protein products using MS to further validate their potential for translation. The statistical models used in MS softwares, such as Proteome Discoverer or MAXQUANT, are well-suited for identifying canonical protein products by scanning around 100,000 translatable protein sequences. However, when searching for 2.4 million lncRNA ORFs (of which only a small fraction is likely to be translated), these tools are prone to generating a significant number of false hits. One potential approach was to narrow down the list of translatable ORFs using ribo-seq, but this risked missing out on potential hits for which ribo-seq reads were missing. To overcome these limitations, we devised a novel strategy by dividing the entire list of ORFs into 30 smaller subsets and conducting separate searches within the MS spectra for each subset, which helped to control the false discovery rate. Each smaller ORF list was merged with human protein sequences downloaded from the REFSEQ database(66,67) and then searched using MAXQUANT(68) (**Fig. 4A**). The resulting peptide hits were compared against known protein sequences to ensure the uniqueness of the identified amino acid sequences. MS data was obtained using the TMT approach, with two replicates from Nalm6K and E cells. The ORFs corresponding to the 1001 peptide hits from all 30 searches were combined and, for additional validation, re-searched using MAXQUANT. Of these, 806 out of 1001 peptide hits were retained in the second round and were considered true hits, henceforth referred to as translatable ORFs (tORFs).

We next compared the relative expressions of the lncRNA derived peptides between Nalm6K and Nalm6E cells (**Fig. 4B**). Among the 806 peptide hits, 50 showed consistent up or downregulation in both replicates of the two cell types (**Fig. 4B**). A majority of the differentially-expressed translatable lncRNAs derived from protein coding genes. We then estimated the log2 fold change of the detected peptides between Nalm6K and Nalm6E cells and plotted them as function of the ORF length from which they originated (**Fig. S3A**). From this, we observe that most lncRNA tORFs are within 100 amino acids in length with a few reaching nearly 500 amino acids.

Next, we analyzed the physicochemical properties of these tORF sequences including the distribution of different amino acids and secondary structure propensities. Overall, the amino acid distribution in the lncRNA tORFs resemble that of known protein sequences in the REFSEQ database (**Fig. 4C**). However, there is a notable abundance within lncRNA tORFs of hydrophobic amino acids such as Met, Leu and Ile and a depletion of amino acids such as aspartic acid (Asp), alanine (Ala), glycine (Gly) and valine (Val). We also predicted the secondary structure propensities and percentage disorder in these ORF sequences using RaptorX(69) (**Fig. 4D**). While 37% of the tORFs were predicted to be completely disordered, the rest were with varying proportions of helices, beta sheets and disordered loops, along with a small percentage being completely ordered (∼18%). Among the sequences predicted to be ordered, we observed a greater propensity towards helical rather than beta sheet structures (**Fig. 4D**). These results are rather striking because abundance of ordered structures within these tORFs may be indicative of functional significance.

To further investigate the functional significance of the tORFs, we used AlphaFold 2.0 (38) to systematically predict their tertiary structures and identify functional protein domains within these structures using InterPro(70) (**Fig. 4E, S3B**). The predicted structures of lncRNA tORFs included both disordered and folded domains. The majority of the folded structures were combinations of helices and disordered loops, while a few were beta hairpins and sheets. Interestingly, we found that 31 tORFs could produce tertiary folds that resembled mini proteins (**Fig. 4E, Fig. S3B**). The ensemble of structures included familiar folds such as helical bundles, beta sheets and a combination of alpha and beta. Also, many of the tertiary folds were stabilized by disulfide bridges, akin to real proteins. Using InterPro, we also found that many of the folded tORFs included functional protein domains that could be broadly classified into RNA/DNA binding and transport/cytoskeletal/enzyme/immune related domains (**Fig. S3B**).

### *SF3B1* mutation associated translatable peptides could serve as neoantigen

Cancer cells are known to produce neoantigens originating from aberrant protein products that can modulate the host immune-landscape or act as targets for immunotherapy(71). To determine whether the tORF-derived peptides could act as neoantigens, we predicted their immunogenicity against several common class-I HLA variants (HLA-A02:01, HLA-B58:01, HLA-C*07:02) using the machine-learning-based program DeepImmuno(40). HLA-A02:01 is one of the most common HLA variants expressed by many individuals, including cancer patients, while HLA-B58:01 and HLA-C*07:02 are associated with various skin diseases. We identified 4,095 peptide epitopes (each 10 amino acids in length) from the 806 tORFs, with immunogenicity scores greater than 0.8, indicating strong reactivity towards the tested HLA variants. Additionally, 449 epitopes were predicted to have strong immunogenicity (score > 0.8) across all three HLA variants, suggesting that these epitopes could play a crucial role in eliciting a host immune response. Notably, 94% of the tORFs contained at least one epitope with strong immunogenicity. Furthermore, several upregulated tORFs in K700E cells exhibited multiple epitopes with strong immunogenicity, implying that these epitopes may be amplified by the *SF3B1* mutation (**Fig. 4F**). Taken together, the diverse tORFs derived from lncRNAs with their potential immunogenicity and ability to fold into functional protein-like structures could be indicative of a hidden proteome within the eukaryotic family and has potentially significant biological functions and disease implications.

### Linear motif enrichments among lncRNA derived peptides and proteins are indicative of functional significance

Given the diversity and folding propensity of the tORFs, we next asked whether the translatable (and foldable) lncRNA-derived protein products could interact with other proteins in the cell and hence influence signaling circuits. Many functional proteins are known to recognize their interaction partners using common sequence motifs referred to as short linear motifs (SLIMs)(72). To explore whether some of these SLIMs are part of the lncRNA-derived tORF sequences, we first investigated their occurrence. We downloaded a set of 300 SLIMs from eukaryotic proteins (eukaryotic linear motif, or ELM) from the ELM resource repository (http://elm.eu.org)(73) and determined the frequencies of these motifs among the 806 tORFs. To assess whether certain ELMs were significantly enriched in differentially expressed tORFs from K700E cells compared to non-differentially expressed ones, we calculated the frequency of each ELM in a set of 806 randomly generated amino acid sequences. These random sequences had identical lengths and similar amino acid distributions to the original lncRNA tORFs. We repeated this process 100,000 times to generate a bootstrapped null distribution that was then used to estimate the statistical significance (p-value) of ELM enrichment. We found that the ELMs were significantly enriched among the upregulated, downregulated and similarly expressed tORFs in the K700E cells (**Fig. S4A-B**). The annotated functions of these ELMs include proteins involved in DNA damage repair and cell cycle regulation, TRAF6 binding motif involved in ubiquitin ligase recognition and NF-kB pathways along with a few of the ELMs representing known phosphorylation motifs found in kinases (**Fig. S4C**). Taken together, these observations suggest that some of the lncRNA-derived proteins upregulated in *SF3B1* mutant cells may modulate major cellular processes such as DNA damage repair and ubiquitination by interacting with key proteins in these pathways. Of note, ELMs are typically short and the risk of finding false positives among such short sequence motifs cannot be discounted. Further investigation is needed to analyze the functional implications of the lncRNA-derived proteome.

### lncRNA tORFs could be detected in CLL leukemia samples with disease implication

To evaluate the disease relevance of the lncRNA-derived tORFs, we investigated their presence in mass spectrometry data from 117 CLL patient samples, obtained through data-independent acquisition (DIA-MS), both with and without the *SF3B1* mutation(32). We found 47 peptide hits from distinct tORFs expressed among multiple patients (**Fig. 5A**). Univariate regression analysis with known CLL molecular and clinical features revealed that deletions in chromosome 11q (*del*(11q), 17p (*del*(17p), and mutations in *ATM, MED12, EGR2, BRAF* and *SF3B1* all with a hazard ratio (HR) greater than 1 (p: 0.01-0.25). Immunoglobulin heavy chain variable region (IGHV) mutation status was also significantly correlated Using the same approach, we identified tORFs whose expression was correlated with time to treatment, such as one tORF from the *ARHGAP12* gene (**Fig. 5C**). Additionally, we identified tORFs whose expression appeared beneficial to the patients, such as one from the CLASP1 gene (**Fig. 5C**).

**Fig. 5.**
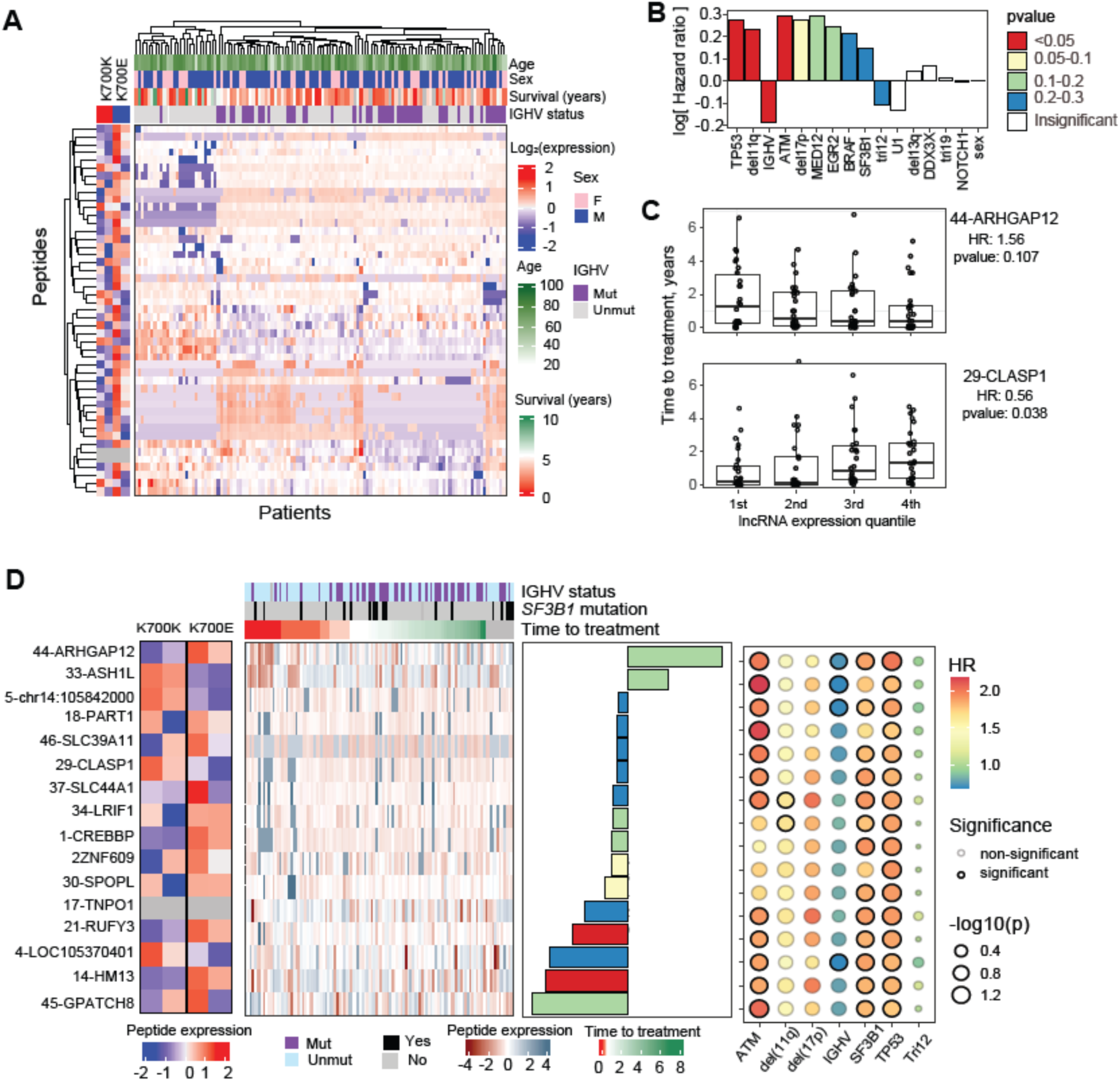
Translated lncRNA peptides have significant clinical associations in chronic lymphocytic leukemia. **(A)** Heatmap showing the quantification of lncRNA derived peptides among CLL patients via DIA mass spectrometry, alongside key clinical attributes (top annotations) and corresponding expressions in Nalm6 samples (left heatmap). **(B)** Top lncRNA derived peptides showing association with time to treatment as identified by univariate regression analysis. bar colors are according to the pvalue; each peptide is labeled by the originating gene name along x-axis. **(C)** Examples of association of lncRNA derived peptides with time to treatment; peptide expression range is divided into quartiles and the median time to treatment in each quartile is presented using boxplots. **(D-E)** Result of multivariate Cox regression of lncRNA derived peptide association with time to treatment, using known CLL risk factors as covariates. Heatmap depicts the expressions of top peptides with significant hazard ratios in individual patients alongside key clinical attributes (top) and corresponding expressions in Nalm6 (left); barplot on the right shows the estimated hazard ratios (bars colored by significance); dotplot shows the contributions of each CLL covariate, colored by hazard ratios; dot sizes are proportional to p value; significant (p value < 0.1) covariates are highlighted using black borders.

To assess the significance of these findings in relation to known CLL risk factors, we performed multivariate regression analysis on each tORF, incorporating select clinical attributes (*del*(11q), *del*(17p), trisomy 12, IGHV, mutations in *SF3B1*, *TP53*) as covariates. This analysis identified 16 tORFs to be significantly correlated with time to treatment (p < 0.25), with two tORFs (ARHGAP12 and ASH1L) showing positive HR (**Fig. 5D**). Most of these tORF analyses exhibited highly significant HR (p < 0.1) that were associated with known CLL risk factors such as mutations in *ATM, SF3B1* and *TP53*, while a smaller subset was significantly linked to *del*(11q) and IGHV mutation status. Furthermore, many of these tORFs were upregulated in *SF3B1* mutant cells, underscoring the role of *SF3B1* mutation in modulating both their expression and disease association. Of note, the top hit, ARHGAP12, showed increased expression in Nalm6 K700E cells and was significantly correlated with IGHV mutation status, as well as mutations in *ATM, SF3B1* and *TP53*. Additionally, this peptide was predicted to have strong immunogenicity (score = 0.92) according to DeepImmuno, and AlphaFold predictions indicate that it adopts an alpha-helical structure with a highly hydrophobic surface, potentially facilitating binding to other proteins and the plasma membrane. Taken together, our *in vitro* model of *SF3B1* mutation combined with long-read sequencing and mass spectrometry has identified protein products derived from translatable lncRNAs that were validated in CLL patient samples, highlighting their potential disease implications and future promises as biomarkers and therapeutic targets.

### Knockdown of tORFs leads to impaired cellular growth in a *SF3B1* mutation-dependent manner

Given the success of our long-read pipeline in detecting large-scale lncRNA translation, we sought to determine whether the resulting protein products could be reproduced in independent experiments using our cell lines. For validation, we selected a tORF from the ZNF609 gene that displayed high conservation across the mammalian family (PhastCons score > 0.9). Notably, the source lncRNA for this tORF was highly expressed in Nalm6K and Nalm6E cells (exceeding 13 transcripts per million reads). Since the ORF is located at the 3’ end of the lncRNA, we fused a FLAG tag to the peptide’s C-terminal end and expressed it in Nalm6K and Nalm6E cells via lentiviral transduction (**Fig. 6A).** We successfully detected tORF protein expression in both cells (**Fig. 6B**), confirming its translation.

**Fig. 6.**
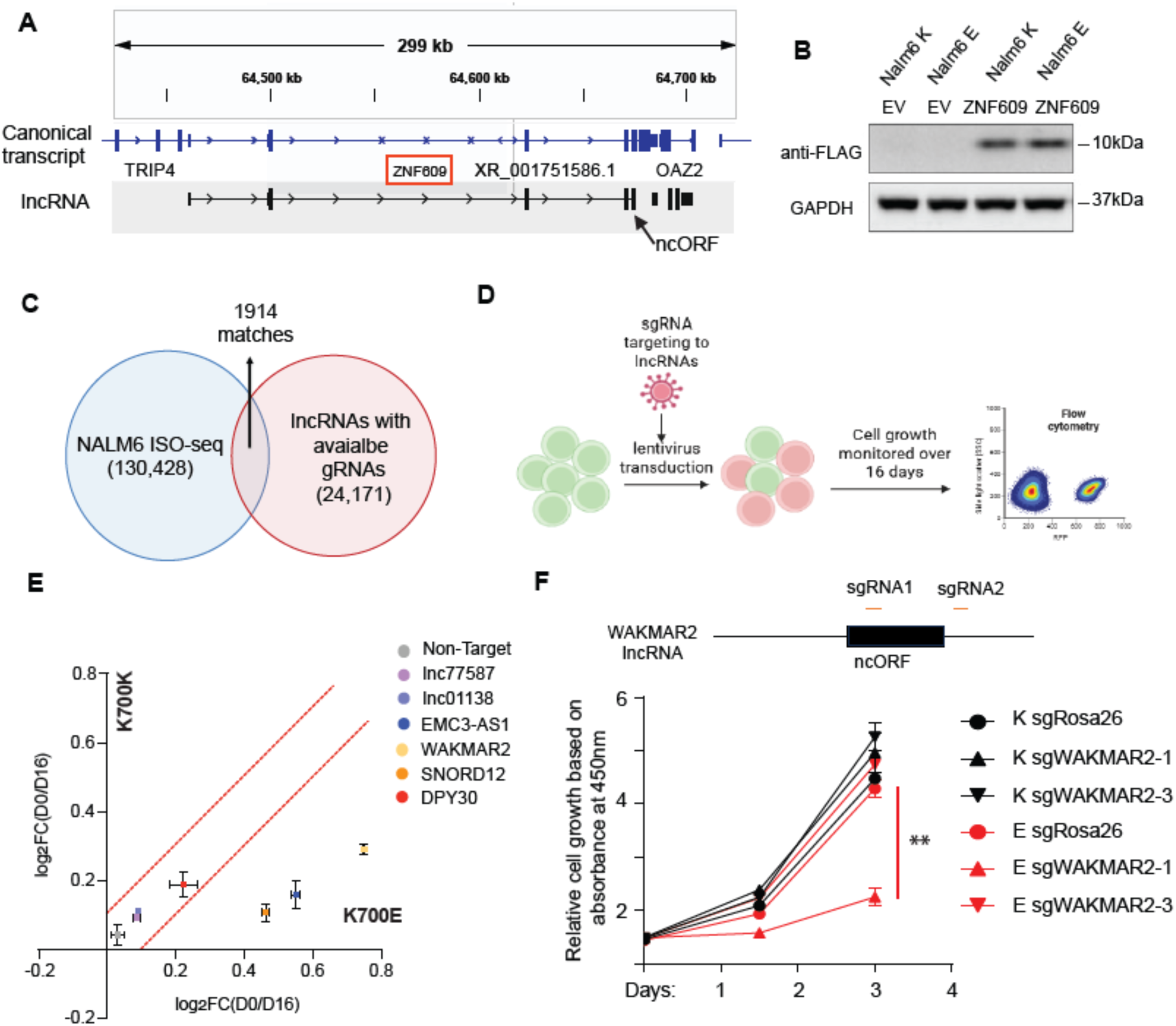
Experimental validation of lncRNA translation and functional significance. **(A)** Genomic location and exonic structure of a translatable lncRNA from the gene ZNF609 shown using Integrative Genomics Viewer (IGV); location of the ncORF is highlighted with arrow. **(B)** Protein product from ZNF609 ncORF translation in Nalm6 cells with and without *SF3B1* mutation from flag-tagged overexpressed ncORF. **(C)** Overlap between lncRNAs from ISO-seq and those with available CRISPR-CasRx gRNAs from Montero et al. **(D)** Workflow for monitoring the effect of CRISPR mediated lncRNA knockdown on cell growth; Nalm6 cells with or without *SF3B1* mutation were transduced with lentivirus targeting lncRNAs and cells were monitored for percentage of RFP by flow cytometry over 16 days. **(E)** Cell growth reduction due to lncRNA knockdown represented as log-ratio of RFP percentage from day 0 over day 16, with higher numbers indicating increased growth reduction; results from wild type and *SF3B1* mutated cells are plotted against each other. **(F)** WAKMAR2 lncRNA schematic showing the two CRISPR-Cas9 gRNA constructs, one within and the other outside ncORF region (upper panel). Relative cell growth based on absorbance at 450 nm from CCK8 assay was plotted against time for the different CRISPR induced Nalm6 cells; error bars are from three replicate measures. ** indicates p<0.01, Student t test.

We next investigated whether the expression of lncRNA-derived tORFs exerts functional effects. A recent study surveyed transcriptome-wide functional lncRNAs using RNA-targeting CRISPR-RfxCas13d (CasRx) screen with 24,171 gRNAs and identified 778 lncRNAs with context-specific or broad essentiality(74). By intersecting this gRNA library with our ISO-Seq data, we identified 1,914 gRNA hits corresponding to 1,778 unique lncRNA isoforms (**Fig. 6C**). In parallel, we assessed the expression of the ISO-Seq-derived lncRNAs in two CLL cell lines (HG3 and MEC1), both with and without the *SF3B1* mutation, using short-read RNA-seq(15) (**Fig. S5A-B**). Encouragingly, 83% of the ISO-Seq lncRNAs from Nalm6 cells (including 683 translatable ones) were also detectable in both CLL cell lines, confirming that the Nalm6 model accurately represents the lncRNA transcriptome of CLL cells (**Fig. S5B**). Moreover, 183 translatable lncRNAs upregulated in Nalm6E were also upregulated in at least one CLL cell line harboring the *SF3B1* mutation; among these, 21 lncRNAs had available gRNAs (**Fig. S5C-D**). Importantly, we identified 11 translatable lncRNAs with corresponding gRNAs.

We selected six highly expressed candidates (*lnc77587, lnc01138, EMC3-AS1, WAKMAR2, SNORD12, DPY30*) from each group for *in vitro* knockdown using CRISPR-CasRx(75), a cutting-edge technology that targets specific RNA transcripts without affecting the parental gene expression (**Fig. 6D-E**). Knockdown of three of the lncRNAs (*WAKMAR2*, *EMC3-AS1*, *SNORD12)* significantly impacted the growth of *SF3B1* mutated cells but not the wild type cells, indicating that their functional effect is *SF3B1* mutation-dependent (**Fig. 6E**). Among these, a translatable lncRNA from the *WAKMAR2* gene induced the most pronounced growth defect. To determine whether lncRNA translation contributes to this effect, we designed CRISPR-Cas9 gRNAs targeting *WAKMAR2* at two distinct locations: one within the tORF region and another outside of it (**Fig. 6F**). We observed significant growth inhibition only in *SF3B1* mutant cells with disruption of the tORF region based on CCK8 cell growth assay (**Fig. 6G**). In contrast, targeting *WAKMAR2* outside the tORF region or knocking out the tORF in Nalm6K cells had minimal impact on cellular proliferation. These findings suggest that the functional effect of *WAKMAR2* is mediated specifically through its tORF region, highlighting the importance of lncRNA translation in the *SF3B1* mutant genomic context.

## DISCUSSION

In this study, we presented a workflow to identify and annotate lncRNAs in the context of splicing factor *SF3B1* mutation with long read sequencing. We identified 83,000 novel lncRNAs associated with *SF3B1* mutation, 806 of them can be translated and detected as peptide from proteomics data. Importantly, we validated the functional impact of translatable lncRNAs working in *SF3B1* mutation-dependent manner. Our study has a few implications for both cancer biology and lncRNA research:

### ALLR: a new workflow to identify and annotate lncRNAs

A longstanding bottleneck in lncRNA studies is the poor functional annotation of large number of lncRNAs. There are a few caveats for the limited knowledge in this space. First, most of the identified lncRNAs were initially identified through a mixture of FANTOM cDNA sequencing efforts and joined by long intergenic noncoding RNA sets, which are considered more conservative collected lncRNAs(1,2). This list was significantly expanded by the incorporation of short-read RNA-seq data sets, which relies on the transcription-reconstruction software to annotate lncRNAs but with limited manual annotation. Hence, the quality is often doubtful. Second, due to the low expression nature of lncRNAs, the coverage of lncRNAs from sequencing studies tends to be low. This has already resulted in a large discrepancy between automated annotation (e.g. 173,112 genes for NONCODE) and highly curated collection (e.g. 53,000 genes for GENCODE). Lastly, the lack of tools to systematically “annotate” lncRNAs *in silico* making it challenging to fully appreciate the biological roles of lncRNAs. Long-read sequencing has revolutionized the annotation in a high throughput fashion as exemplified from a recent publication, in which capture long-read sequencing was utilized to annotate full-length lncRNAs(6). Motivated by this approach, we applied long-read sequencing technology to identify splicing factor mutation associated lncRNAs. Moreover, leveraging available short-read RNA-seq and proteomic dataset derived from the same cells, we developed a powerful workflow to annotate the potential functions of lncRNAs. This led us to discover disease-associated lncRNAs as well as a large set of translatable lncRNAs. We are confident that our approach could help to discovery key lncRNAs with physiological and pathophysiological roles in normal process and disease settings.

### Splicing factor mutation influences landscape of lncRNAs and may employ lncRNAs to impact cancer biology

Worldwide cancer genome sequencing efforts have identified high frequency mutations of spliceosome components (i.e. *SF3B1, U2AF1, SRSF2, ZRSR2,* U1snRNA) in multiple hematological malignancies and solid tumors(8,10,76,77). In particular, *SF3B1* mutation was recurrent mutated across different cancer types with strong clinical outcome correlation(8–10,14,77). We and others have consistently found that *SF3B1* mutation impacts mRNA splicing with a preferential influence on the selection of 3’ ss selection and intron retention(13,23,55,56). While previous studies are restricted to mRNA splicing, here, we explore the role of this cancer-driving mutation in the biogenesis of lncRNA. Our results clearly demonstrate that *SF3B1* mutation has an essential role in the generation of lncRNAs as well as the splicing of lncRNAs, with a similar rule followed as mRNA splicing. Importantly, our workflow provided critical information regarding the potential functions of these newly identified lncRNAs, at least one of them through translation.

Our discovery of hundreds of translatable ORFs, many of which with predicted folded structures, enriched functional motifs, and detectable expression in primary CLL samples, reveals the existence of a mutation-dependent “hidden proteome”. Importantly, the strong predicted immunogenicity of these peptides, including multiple high-affinity epitopes across common HLA alleles, positions splicing factor mutation-derived lncRNA peptides as an underexplored, tumor-specific source of neoantigens. Because these epitopes arise from aberrant splicing events unique to mutant cells, they may provide improved specificity for immunotherapeutic targeting compared to conventional mutation-derived neoantigens. Indeed, *SF3B1* mutation-driven neoepitopes have been experimentally proven in uveal melanoma, highlighting the potential for identifying shared neoantigen across different cancer types. Our data provides a foundation for such efforts, extending beyond *SF3B1* to other splicing factor mutations.

Beyond their immunologic relevance, our work also highlights lncRNAs as functional drivers of mutation-dependent cancer growth. CRISPR-CasRx disruption of specific tORFs impairing proliferation selectively in *SF3B1* mutant cells demonstrates that lncRNA translation is biologically consequential and genetically context dependent. Thus, splicing factor mutations reshape the lncRNA landscape in two synergistic ways: by expanding the pool of immunogenic peptides and by generating novel growth-supportive microproteins that reinforce malignant fitness, offering new avenues for therapeutic targeting and biomarker discovery.

In summary, ALLR provides a scalable and systematic framework to uncover multiple layers of lncRNA regulation, establishing *SF3B1* mutation associated translatable lncRNAs as a promising new frontier for biomarker discovery, mechanistic investigation, and therapeutic targeting in spliceosome-mutant malignancies. By revealing translatable lncRNAs and their potential functional peptides, our work offers novel insights into the molecular consequences of *SF3B1* mutations and their role in shaping disease progression, highlighting opportunities to exploit these previously hidden proteomic and regulatory networks for clinical benefit.

## ACKNOWLEDGEMENTS

This work was supported by grants from the National Institutes of Health (NCI) 1R01CA240910, R01CA282450 and R01CA283171 (to L.W.). Work performed in the Integrative Genomics Core at City of Hope (S.B.) supported by NCI grant P30CA033572 (institutional grant). The content is solely the responsibility of the authors and does not necessarily represent the official views of the NIH.

## AUTHOR CONTRIBUTIONS

Supriyo Bhattacharya: Conceptualization, Formal analysis, Methodology, Software, Visualization, Validation, Data curation, Writing—original draft, review & editing. Mike Fernandez: Formal analysis, Methodology, Visualization, Validation, Writing—review & editing. Yiming Wu: Methodology, Writing—review & editing. Meiling Jin: Methodology, Formal analysis, Writing—review & editing. Patrick Pirrotte: Methodology, Formal analysis, Writing— review & editing. Krystine Mansfield: Methodology, Formal analysis, Writing—review & editing. Tae Hyuk Kang: Methodology, Formal analysis, Writing—review & editing. Yuqi Zhao: Methodology, Formal analysis, Writing—review & editing. Nicole Lammana: Methodology, Writing—review & editing. Xiwei Wu: Methodology, Formal analysis, Writing—review & editing. Lili Wang: Conceptualization, Project administration, Supervision, Funding acquisition, Resources, Data curation, Methodology, Writing—original draft, review & editing.

## SUPPLEMENTARY DATA

### Supplementary Data statement

Supplementary Data are available at NAR online.

## DATA AVAILABILITY

The ISO-seq and Ribo-seq data from Nalm6 K700K and E are deposited in GEO under accession IDs GSE319973 and GSE319974. The RNA-seq data from Nalm6 and CLL cell lines were published previously (GEO IDs: GSE72790 and GSE231387 respectively). The proteomics data from Nalm6 cell lines are deposited in PRIDE under accession number PXD078356. DIA-MS data from CLL patients were obtained from PRIDE (PRIDE ID: PXD022198). All processed data and scripts needed to reproduce the work in this manuscript have been deposited in Zenodo (10.5281/zenodo.20787494). Additional data and information can be obtained by contacting the corresponding author.

