## Supplementary figures and tables for "*SF3B1* mutation generates novel peptides from long non-coding RNAs"

**MANUSCRIPT TITLE**

**Supplemental Figures**

(A)

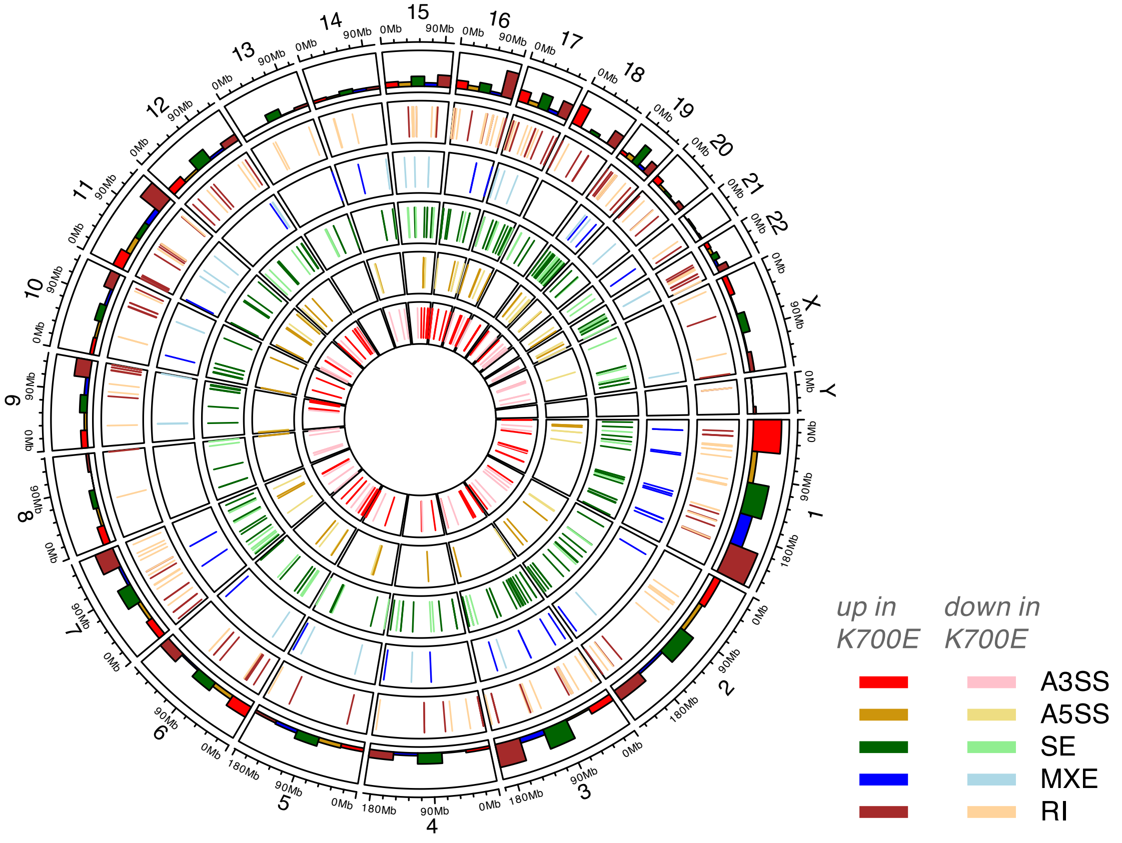


(B)

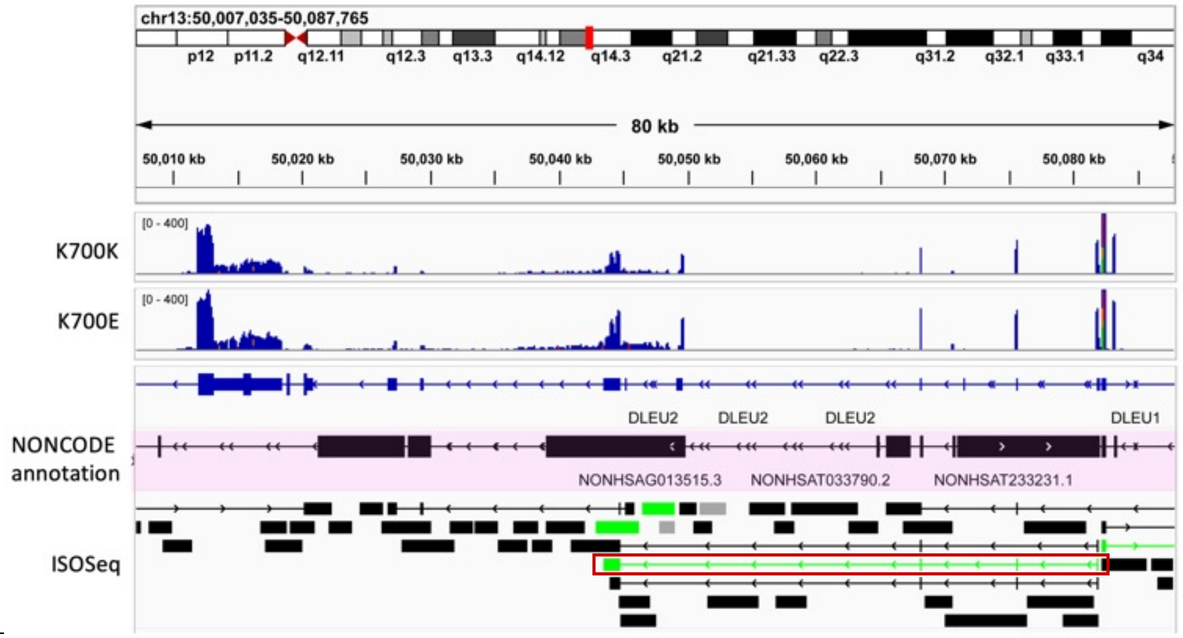

**Figure S1. ISOseq detects differentially expressed alternative splicing events and novel lncRNAs due to K700E mutation.** (A) Chromosome-wise distribution of alternative splicing events (color-coded according to category) within the human genome, that are differentially expressed between K700K and E. Bar plots on the outermost sector represent the total number of events per chromosome detected in each category. (B) IGV snapshot showing the chromosomal region corresponding to the upregulated lncRNAs that are implicated in CLL. The first two tracks represent the RNASeq read counts in K700K and K700E. The track highlighted in pink shows the location of the annotated lncRNAs from the NONCODE database. The ISOSeq track shows the different lncRNA isoforms that overlap with the DLEU2 region in the genome. Upregulated lncRNAs in K700E are highlighted in green. (B) long-read sequencing (ISO-seq) identified a novel DLEU2 lncRNA. Genome browser snapshot of the 13q14.3 region (chr13:50,007,035–50,087,765), encompassing ~80 kb. Cytoband and scale bar indicating chromosomal position and direction on the top. RNA-seq coverage from Nalm6 K700K and K700E cells, demonstrating differential expression of DLEU2 across the locus. NONCODE annotation, highlighting known lncRNAs including DLEU1, DLEU2, and sub-isoforms (pink). ISO-seq data reveal multiple novel lncRNA isoforms, including a transcript (highlighted in red box) not annotated in existing databases but supported by full-length reads (bottom).

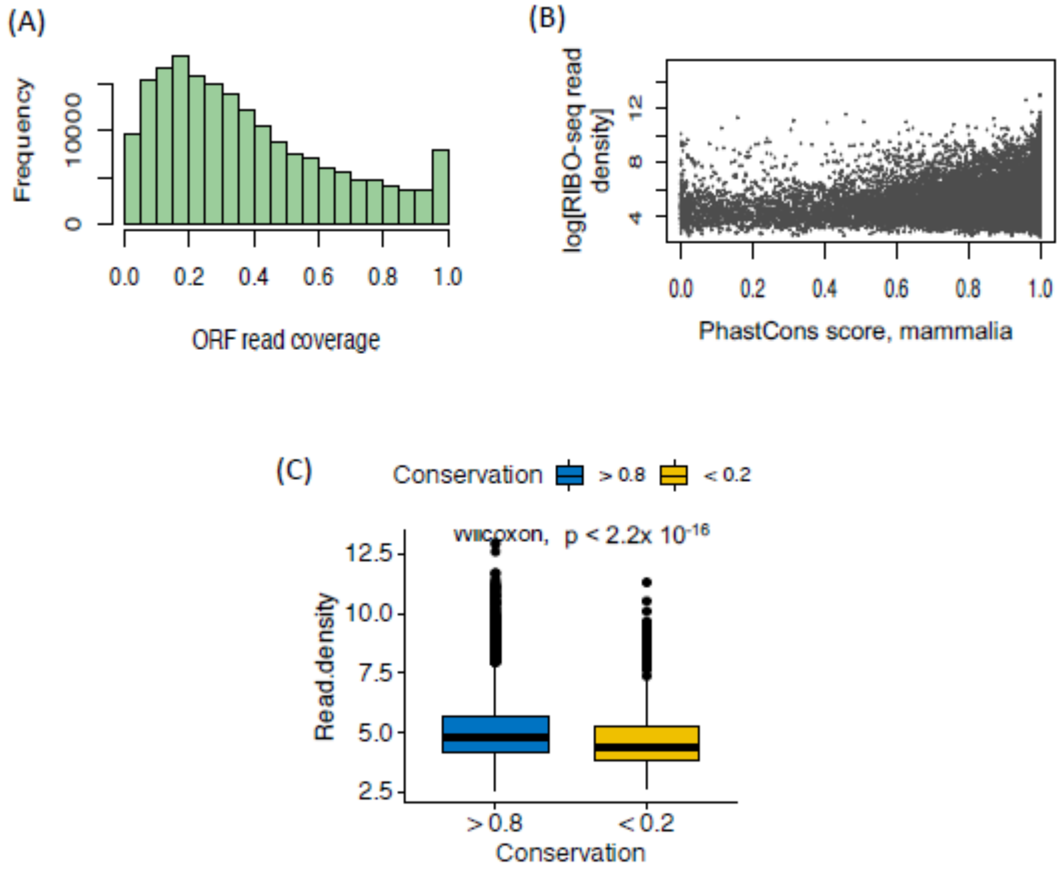


**Figure S2. Association between ORF translation, evolutionary conservation, and ribosome occupancy of *SF3B1* mutation associated lncRNAs. (A)** Histogram of ORF read coverage across candidate open reading frames (ORFs) shows a broad distribution of ribosome footprint coverage with a peak around 0.3. **(B)** Scatter plot showing a positive trend between mammalian PhastCons conservation scores (x-axis) and ribosome profiling (Ribo-seq) read density (log scale, y-axis). More conserved sequences tend to have higher ribosome occupancy. **(C)** Box plot comparing Ribo-seq read densities of highly conserved (PhastCons > 0.8, blue) versus poorly conserved (PhastCons < 0.2, yellow) ORFs. Significantly higher ribosome occupancy is observed in conserved ORFs (Wilcoxon test, $p < 2.2 \times 10^{-16}$).

(A)

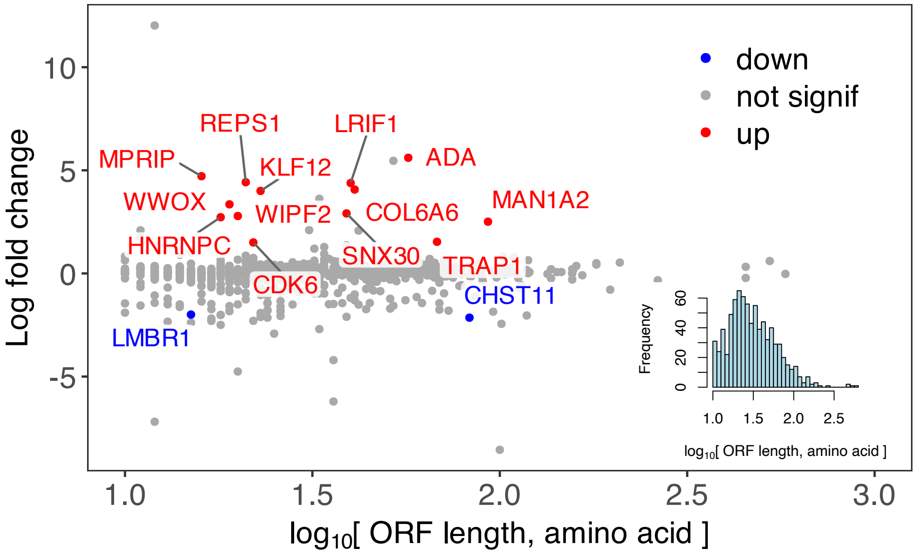


(B)

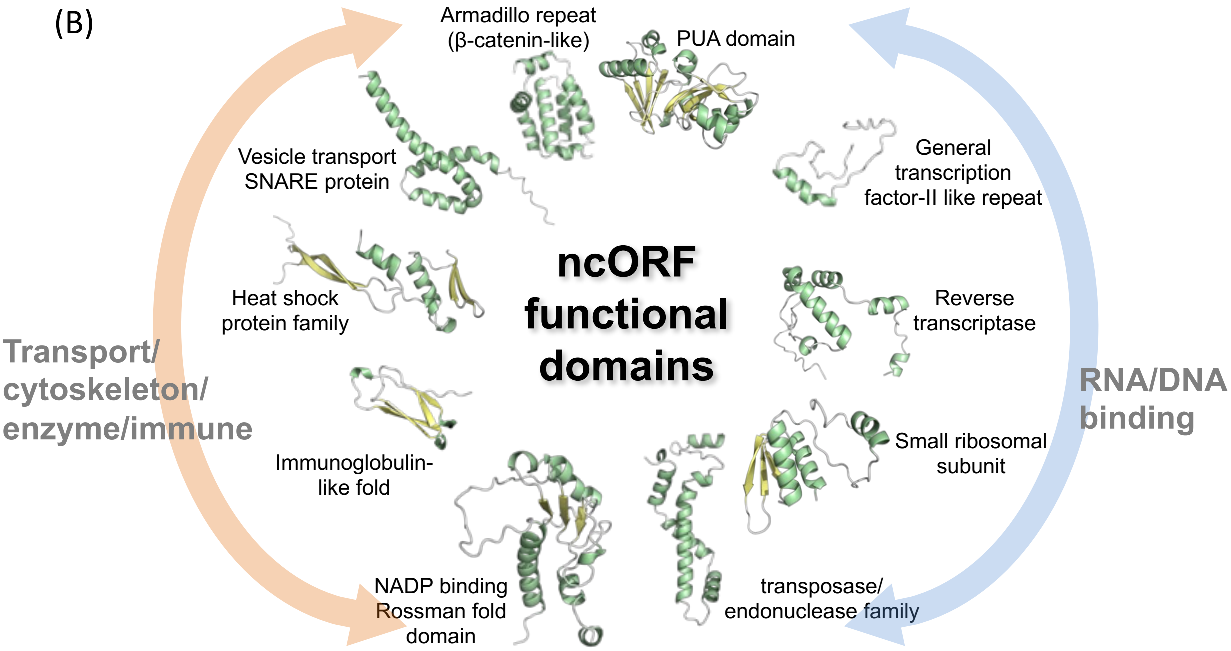


**Figure S3. Differential expression and predicted functional domains of non-canonical ORFs (ncORFs). (A)** Plot of ORF length vs log fold change (E vs K) of the detected peptides. Peptides with LFC > ±2 are highlighted in red (up in K700E) and blue (down in K700E). Inset: Frequency distribution of ORF length.Inset: Histogram showing the distribution of ORF lengths ($log_{10}$ scale), highlighting the prevalence of short ncORFs. **(B)** Predicted protein domains identified among ncORFs using structural modeling and functional annotation.

(A)

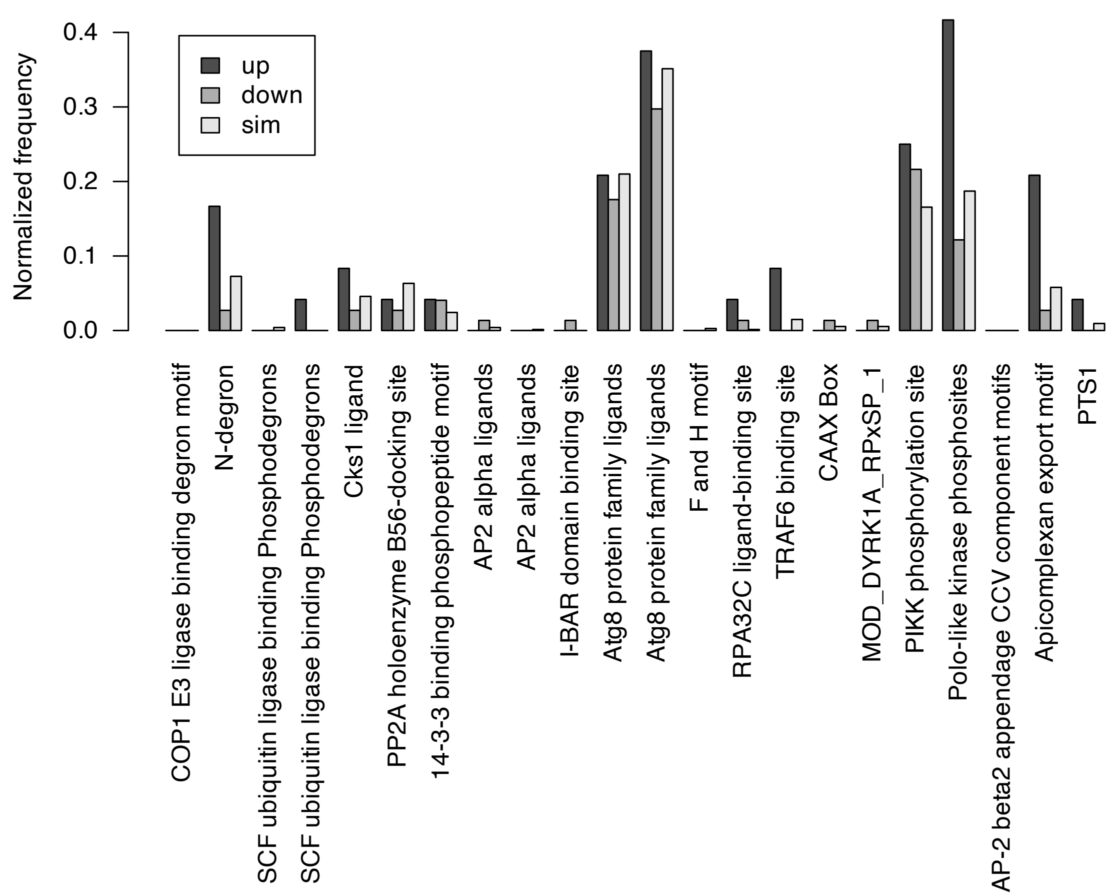


(B)

**TRAF6 binding motif:** PxExx[F/Y/W/H/D/E]
lncRNA gene: MAN1A2
...KKSTYSQDNPKPKEQSWRHHIT

**SCF ubiquitin ligase binding phosphodegron:** [L/I/V/M/P]xTPxxE
lncRNA gene: KIAA0753
...IFFNGYLAIPTPSDESNTWTVC

**PIKK phosphorylation site:** xxx[S/T]Qxx
lncRNA gene: FTO
...NQYGTLKPSCNSQEKTILAFKIS

(C)

| lncRNA peptide ELM | Annotated function |
|---|---|
| Apicomplexan export motif | Exports secreted proteins from infecting protozoa into cytosol |
| PIKK phosphorylation site | DNA damage repair |
| N-degron | N-terminal protein degradation motif |
| Cks1 ligand | Binding motifs for CDKs, E3 ligases etc. (involved in cell cycle and cell growth) |
| TRAF6 binding site | E3 ubiquitin ligase and NF-kB pathways |
| Atg8 protein family ligands | Autophagy pathways |
| RPA32C ligand-binding site | DNA damage repair |
| SCF ubiquitin ligase binding Phosphodegrons | Phosphorylation dependent ubiquitin ligase binding |
| Polo-like kinase phosphosite | Cell cycle, mitosis, DNA damage response |

**Figure S4. Enrichment and functional annotation of peptide motifs in translated lncRNA-encoded ncORFs. (A)** Bar plot showing the normalized frequency of Eukaryotic Linear Motifs (ELMs) in upregulated (dark gray), downregulated (black), and similarly expressed (light gray) lncRNA-derived peptides. Several motifs (e.g., TRAF6 binding site, WD40 binding ligand, SCF ubiquitin ligase site) are significantly enriched in the upregulated group ($p < 0.05$, asterisks). **(B)** Representative examples of enriched peptide motifs detected in upregulated lncRNAs. Red letters highlight motif cores in TRAF6 binding motif, SCF ubiquitin ligase binding phosphodegron, and CDK phosphorylation site. **(C)** Table summarizing selected enriched ELMs, corresponding peptide motifs, and their annotated biological functions. These motifs suggest potential roles of lncRNA-encoded peptides in protein–protein interaction networks, post-translational regulation, and key signaling pathways (e.g., ubiquitination, DNA damage repair, cell cycle control).

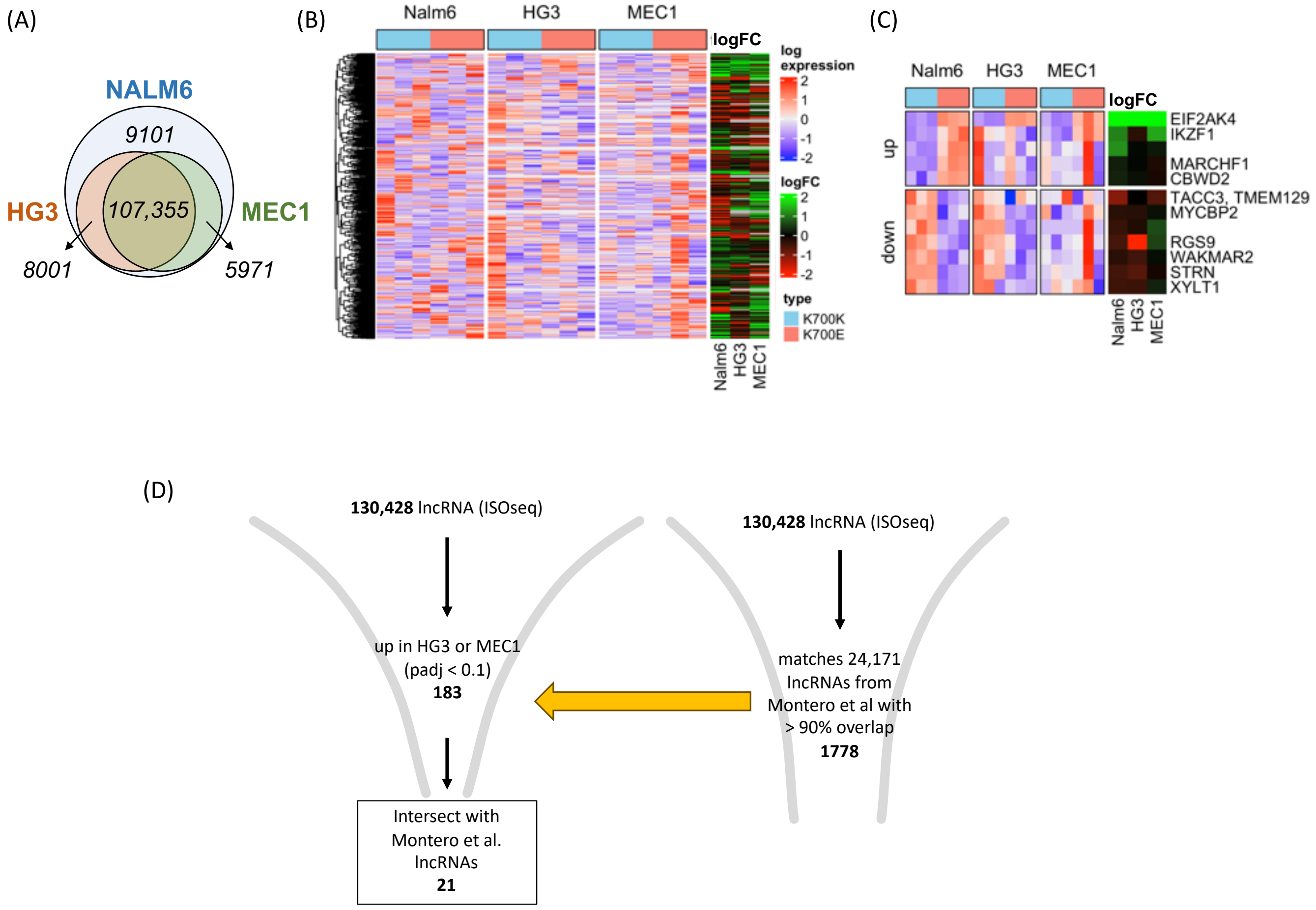

**Figure S5. Experimental validation of candidate lncRNAs identified from ISO-seq and differential expression analysis across leukemia cell lines. (A)** Venn diagram showing overlap of lncRNAs detected by ISO-seq in three cell lines: NALM6 (blue), HG3 (orange), and MEC1 (green). A core set of 107,355 lncRNAs is shared among all three. **(B)** Heatmap of translatable lncRNA expression profiles and log fold change (logFC) across NALM6, HG3, and MEC1. Differential expression is shown for both K700K and K700E isoforms. **(C)** Heatmap of representative translatable lncRNAs showing consistent upregulation or downregulation in HG3 and MEC1 compared to NALM6. Gene names were labeled. (D) Strategy for identification of SF3B1 mutation associated key functional lncRNAs. Out of 130,428 lncRNAs from ISO-seq, 1,778 lncRNAs matched Montero's dataset with >90% overlap among 24,171 sgRNA-available lncRNAs, 183 were upregulated in HG3 or MEC1 (padj < 0.1). 21 overlapped with lncRNAs reported by Montero et al. These were selected for further functional validation.

**Supplementary table S1.** Number of RNA isoforms from different categories retrieved by $ISOseq^{lowres}$ and $ISOseq^{hires}$

| | **$ISOseq^{lowres}$** | **$ISOseq^{highres}$** | **Common** |
|---|---|---|---|
| **Protein coding** | 52,386 | 72,296 | 50,649 |
| **Noncoding** | 73,082 | 130,428 | 7,920 |
| **Total** | 125,468 | 202,724 | 58,569 |

**Supplementary table S2.** Number and quantification of splicing events in different categories identified using $ISOseq^{hires}$ derived splicing junctions

| **Event type** | **Detected events** | **Differentially spliced, K700E vs K** | **% differentially spliced** |
|---|---|---|---|
| **A3SS** | 9149 | 192 | 2.1 |
| **A5SS** | 9172 | 79 | 0.86 |
| **SE** | 38178 | 259 | 0.68 |
| **MXE** | 9703 | 76 | 0.78 |
| **RI** | 33048 | 309 | 0.94 |

**Supplementary table S3.** Number and quantification of splicing events in different categories identified using GENCODE annotated splicing junctions

| Event type | Detected events | Differentially spliced, K700E vs K | % differentially spliced |
|---|---|---|---|
| **A3SS** | 7885 | 113 | 1.43 |
| **A5SS** | 4970 | 26 | 0.52 |
| **SE** | 27411 | 214 | 0.78 |
| **MXE** | 5413 | 51 | 0.94 |
| **RI** | 4217 | 52 | 1.23 |

**Supplementary table S4.** Number and quantification of splicing events in different categories identified using ISOseq$^{lowres}$ derived splicing junctions.

| Event type | Detected events | Differentially spliced, K700E vs K | % differentially spliced |
|---|---|---|---|
| **A3SS** | 4710 | 118 | 2.51 |
| **A5SS** | 4430 | 47 | 1.06 |
| **SE** | 21138 | 174 | 0.82 |
| **MXE** | 5754 | 35 | 0.61 |
| **RI** | 11687 | 101 | 0.86 |

**Supplementary table S5.** Cloning primer sequences for ZNF609.

| Name | Sequence |
|---|---|
| 3xFLAG-ZNF609-SpeI-F | aaaaACTAGTATGgactacaaagaccatg |
| 3xFLAG-ZNF609-BamHI-R | aaaaGGATCCCTACAAGTCGAACT |

**Supplementary Table S6**. Guide RNA sequences for CRISPR/Cas9 and CasRx targeting.

| Name | Nuclease | Sequence |
|---|---|---|
| WAKMAR2 sgRNA1 | Cas9 | ATCCACGGAGTCACAAAAAC |
| WAKMAR2 sgRNA2 | Cas9 | GATAAGAAGGCCAATACCCA |
| WAKMAR2 gRNA1 | CasRx | AAAAGTATTGCAAATCCCCAGAGCCCCATT |
| WAKMAR2 gRNA2 | CasRx | ACAGCCCAAACTTTTCAGAGCCGGCCCGAG |

| | | |
|---|---|---|
| EMC3-AS1 gRNA1 | CasRx | ACATGCCACAAAAAGCTGTCAGCATCTGCC |
| EMC3-AS1 gRNA2 | CasRx | GTAACAATATTGACCCCACCAACTTGATTT |
| LINC01138 gRNA1 | CasRx | ATAAGAGAAATTCGAGAAGGCAATGGGGCA |
| lnc77587 gRNA1 | CasRx | CCCCATATATCTTCACCACTCCAGGTCTCA |
| SNORD12 gRNA1 | CasRx | CATACAAAGTTAAAGACCCCAGGGGAAATA |
| DPY30 gRNA1 | CasRx | ACATTACTATCAGTAGGACCACGGTTTGGA |

**Supplementary Table S7**. Quantitative PCR primer sequences

| **Name** | **Sequence** |
|---|---|
| ZNF609 mRNA F | AGAAAAAGAAAGACAAAAAAAAGAAGG |
| ZNF609 mRNA R | TCTCCTGAAGATTCCCTGAATG |
| lncZNF609 F | GGGTGACTCTGGTTGTCC |
| lncZNF609 R | GGACAACATCATTGCGCG |
| GAPDH F | GGAGCGAGATCCCTCCAAAAT |
| GAPDH R | GGCTGTTGTCATACTTCTCATGG |
